# Enhancing glymphatic function with very low-intensity ultrasound via the transient receptor potential vanilloid-4-aquaporin-4 pathway

**DOI:** 10.1101/2023.01.13.523878

**Authors:** Wei-Hao Liao, Chueh-Hung Wu, Ya-Cherng Chu, Ming-Yen Hsiao, Yi Kung, Jaw-Lin Wang, Wen-Shiang Chen

## Abstract

Recently, the glymphatic system has been proposed as a mechanism for waste clearance from the brain parenchyma. Glymphatic dysfunction has been associated with several neurological diseases such as Alzheimer’s disease, traumatic brain injury, and stroke. Therefore, it may be an important target for therapeutic interventions. In this study, we demonstrated that very low intensity ultrasound (VLIUS) (center frequency = 1 MHz; pulse repetition frequency = 1 kHz; duty factor = 1%, and spatial peak temporal average intensity [I_spta_] = 3.68 mW/cm^2^; duration = 5 min) could significantly enhance the influx of cerebrospinal fluid tracers into the perivascular spaces of the brain and also facilitate interstitial substance clearance from the brain parenchyma. Notably, no evidence of brain damage was observed after VLIUS stimulation. We also demonstrated that VLIUS enhanced the glymphatic influx via the transient receptor potential vanilloid-4-aquaporin-4 pathway in the astrocytes. This mechanism may provide insights into VLIUS-regulated glymphatic function that modifies the natural course of central nervous system disorders related to waste clearance dysfunction.

**One Sentence Summary:** Very low-intensity ultrasound enhances glymphatic influx via the TRPV4-AQP4 pathway in the astrocytes, without observable brain damage.

## INTRODUCTION

Waste clearance is important for the central nervous system (CNS) to maintain normal functioning. The concept, termed the “glymphatic system,” has recently been proposed as a mechanism for waste clearance from the brain parenchyma. The current model of the glymphatic system describes the influx of cerebrospinal fluid (CSF) from the subarachnoid space into the brain parenchyma through the periarterial spaces, mixing with parenchymal interstitial fluid (ISF) and waste products facilitated by aquaporin-4 (AQP4) channels in the end-feet of the astrocytes, and drainage through the perivenous spaces(*1-5*). Glymphatic dysfunction has been associated with several neurological diseases, including Alzheimer’s disease(*6, 7*), traumatic brain injury(*5, 8, 9*), stroke(*10, 11*), migraine(*12*), multiple sclerosis(*13*), and amyotrophic lateral sclerosis(*14*). Due to its pathophysiological association with a broad range of CNS diseases, the glymphatic system may be an important target for therapeutic interventions.

Interactions between the glymphatic system and various intrinsic and extrinsic factors, such as sleep, body posture, blood pressure, aging, and anesthesia, have been reported. Glymphatic influx decreases following sleep deprivation(*15*), and is more effective in the right-lateral decubitus position than in the prone position(*16*). Epinephrine-induced acute hypertension considerably reduces the influx of CSF tracers(*17*). Decreased glymphatic clearance has been observed in older brains(*18*). Some anesthetics (xylazine, dexmedetomidine) show a higher CSF tracer influx, similar to that observed during spontaneous sleep, while others (pentobarbital, isoflurane) significantly inhibit glymphatic influx(*19, 20*). Since the glymphatic system helps in CNS waste removal, such as amyloid-β and tau proteins, and can be modulated by various factors, exploring a way to enhance glymphatic function may be one of the therapeutic approaches for CNS disorders related to waste clearance dysfunction.

Aquaporin-4 (AQP4), with its paravascular polarization, is related to glymphatic function. AQP4-knockout mice had unperturbed influx within the periarterial spaces, but the tracer flow from these spaces to the surrounding parenchyma was significantly impaired(*1*), suggesting that AQP4 facilitates fluid movement between the paravascular and interstitial spaces. Snta1-knockout mice with regular AQP4 expression but AQP4 polarization loss had reduced glymphatic flow as in the AQP4-knockout mice(*21*). Calmodulin (CaM), which drives AQP4 cell surface localization by binding to the carboxyl terminus of AQP4, can be inhibited by trifluoperazine for reducing brain edema(*22*). A complex containing AQP4 and transient receptor potential vanilloid 4 (TRPV4) in astrocytes is essential for maintaining brain volume homeostasis(*23*). Therefore, modulation of TRPV4/CaM/AQP4 may affect glymphatic function. Mechanical stimulation may affect glymphatic function. One study showed that fluid shear stress, analogous to that produced by paravascular CSF or ISF dynamics, can mechanically stimulate N-methyl-D-aspartate receptors on astrocytes, producing increased calcium ion (Ca^2+^) currents and suggesting mechanotransduction in the glymphatic flow(*24*). Another study showed the crucial role of TRPV4 in ultrasound-mediated blood-brain barrier permeability(*25*).

Transcranial ultrasound has been shown to enhance the influx of cerebrospinal fluid into the perivascular spaces of the brain, glymphatic system, and brain parenchyma(*26*). Moreover, a recent study indicated that ultrasound enhances the glymphatic–lymphatic clearance of Aβ predominantly by increasing brain-to-CSF Aβ drainage(*27*). Based on these findings, we hypothesized that ultrasound has a potential role in the treatment of degenerative CNS disorders, and to be one of the mechanisms that enhances glymphatic function via the TRPV4/AQP4 pathway.

In this study, we investigated the ultrasound-induced alterations in glymphatic dynamics. Our previous study revealed the effect of very low-intensity ultrasound (VLIUS) on inducing neurogenesis in specific regions of a mouse brain without negative effects(*28*). In the current study, we explored the relationships between VLIUS, TRPV4 and AQP4, in glymphatic function modulation.

## RESULTS

### VLIUS increased CSF tracer influx

First, we evaluated the effect of different ultrasound intensities on the circulation of the glymphatic system, and the results showed that the intensity at I_spta_ = 3.68 mW/cm^2^ could effectively promote the diffusion of the tracer in the brain (Supplementary Fig. S2 and Fig. 1**A**). In addition, since we used 500 µm thick tissue combined with the tissue clearing process, we obtained more detailed image results, such as the amount and depth of CSF expansion to the paravascular spaces; and showed the amount and depth of tracer expansion into the paravascular spaces could also be significantly increased by VLIUS stimulation (Fig. 1**B–D**). Measurements of tracer penetrance at various slice positions relative to bregma revealed that VLIUS stimulation increased CSF penetrance throughout the posterior (bregma -2) to anterior (bregma +1) sections, indicating the effect was not region-specific (Fig. 1**E**). Quantification of tracer intensity in the cortical parenchyma indicated depth-dependent profiles decayed with cortical depth after the initial peak in both the VLIUS-stimulated and control mice. Notably, the tracer signals in the cortex of the VLIUS-stimulated mice had a higher intensity and penetrated more deeply into the parenchyma (Fig. 1**F**). In vivo transcranial live imaging of the CSF tracer showed greater CSF influx in the VLIUS stimulation group than in the control group at 15 minutes (Fig. 1**G**, Videos 1 and 2). The tracer was distributed into the brain parenchyma through a network of paravascular spaces in cerebral arteries on the brain surface. More tracer infiltration into the brain was observed in the VLIUS-stimulated mice than in the control mice at all observation times.

**Fig. 1:**
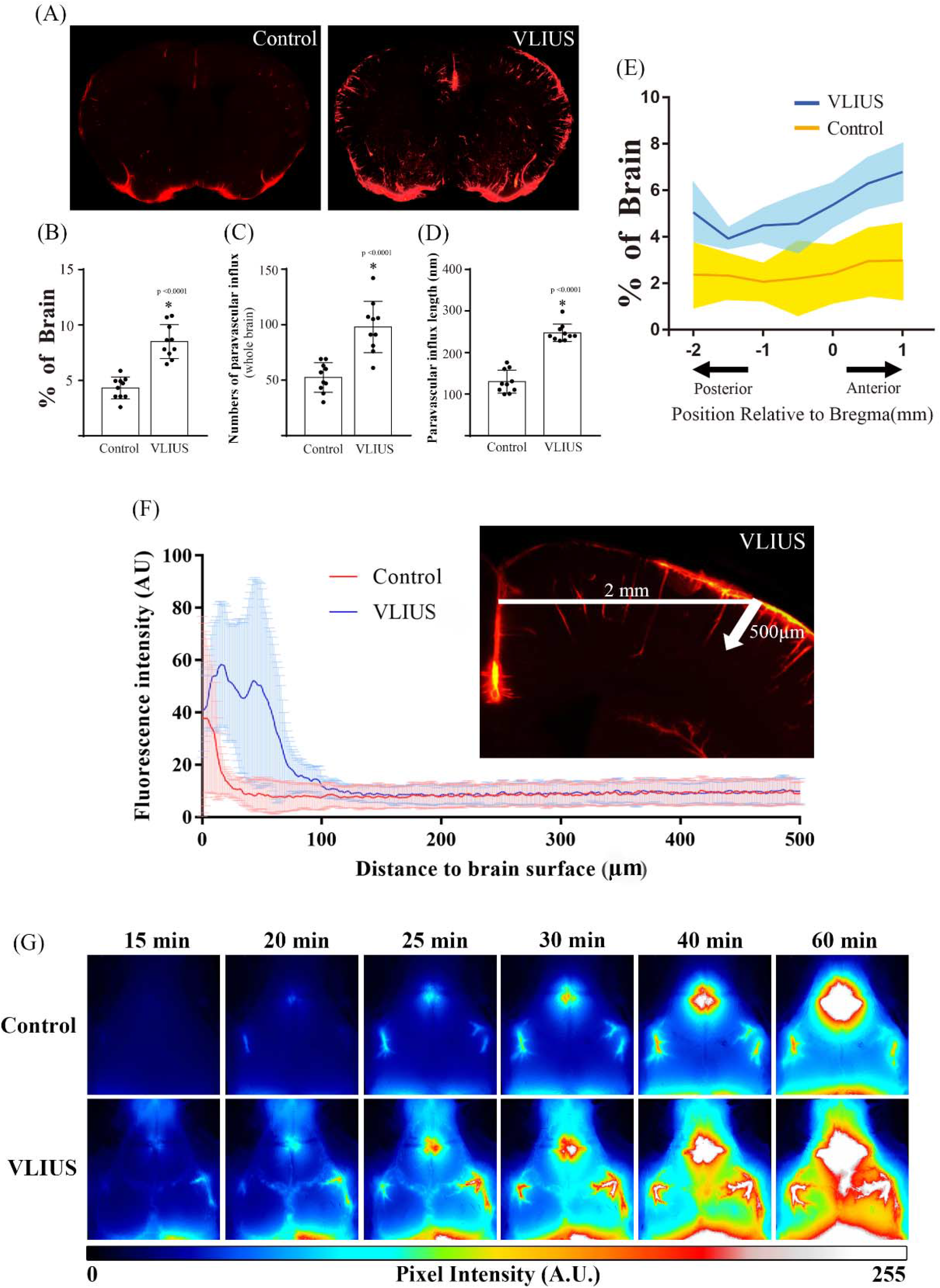
VLIUS stimulation increases CSF tracer penetrance. (**A**) Representative images of coronal brain sections at +0 mm from bregma show increases in CSF penetrance in response to VLIUS as compared to control. Quantification of influx area (**B**), influx numbers (**C**), and influx length (**D**) of VLIUS stimulation compared to controls are shown, with dots representing individual mice in each group. n = 5 mice/group; 2 independent repeats (total n = 10 mice/group); (**E**) Positional slice-by-slice representation of the area covered by tracer influx in coronal brain slices relative to bregma. The solid line represents the percentage of average tracer influx of all brain slices at that section per condition (shaded area = ±STDEV). n = 5 mice/group; 2 independent repeats (total n = 10 mice/group) (**F**) Tracer penetration depth was measured at the cortical position 2 mm lateral to the midline, from the pial surface to a depth of 500 μm in the coronal section. Quantification of the mean fluorescence intensity (MFI) of tracer indicates that VLIUS stimulation induces greater penetration of the tracer deep into the brain (n = 5, respectively). (**G**) Representative time-lapse images of CSF influx over the first 60 minutes following tracer injection in control and VLIUS-stimulated mice. Images (8-bit pixel depth) are color coded (royal form ImageJ) to depict pixel intensity (PI) in arbitrary units (AU). CSF, cerebrospinal fluid; VLIUS, very low intensity ultrasound.

### VLIUS enhanced interstitial substance clearance

In addition to promoting CSF tracer influx, we analyzed whether VLIUS promoted tracer removal from the anterior striatum. Three hours after intrastriatal injection, the mice were euthanized and the tracer residues in their brains were analyzed. VLIUS stimulation significantly reduced tracer residues in the injection area (Fig. 2), implying the possibility that VLIUS promotes waste clearance in the brain.

**Fig. 2:**
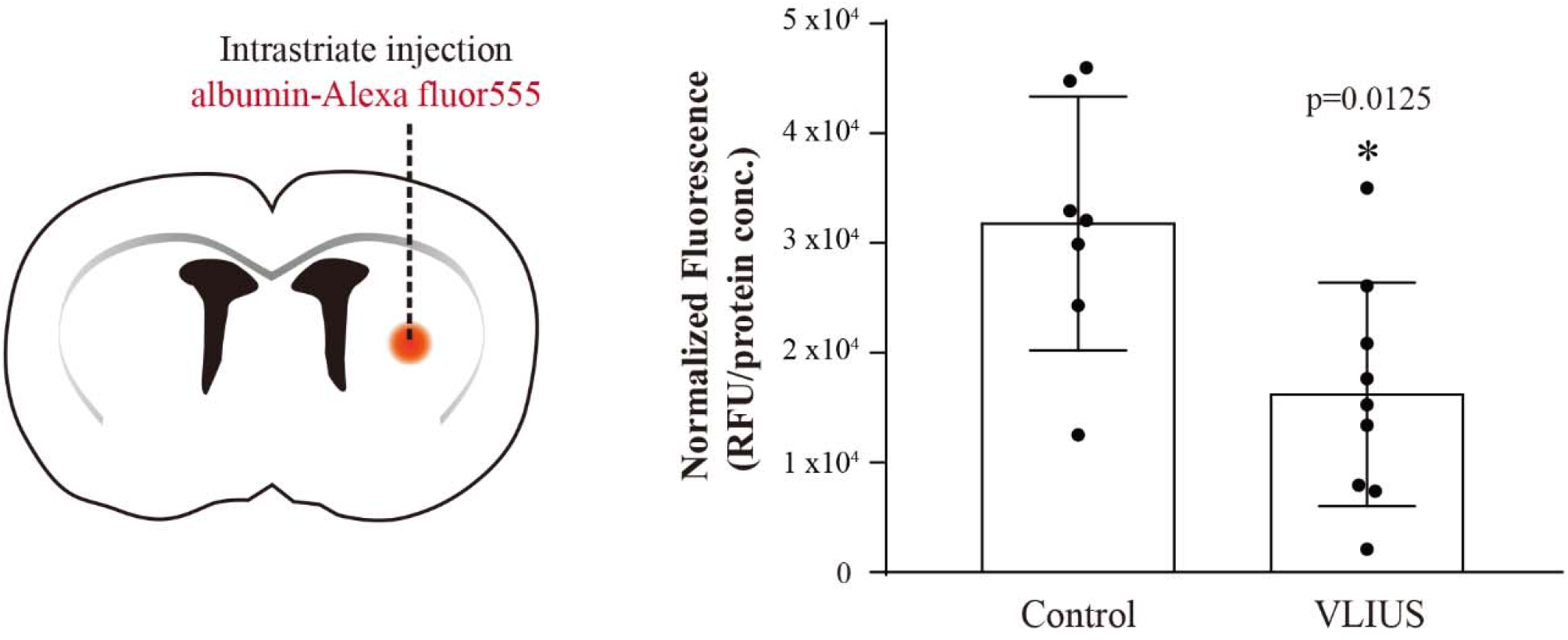
Interstitial fluid clearance greater through VLIUS stimulation. **Left**: Schematic of the experiment. **Right**: Three hours after injection, the remaining tracer was significantly lower in very low intensity ultrasound (VLIUS)-stimulated mouse brains than in the controls (p = 0.0125, control group n = 7, VLIUS group n = 9), indicating that the clearance rate may be higher under VLIUS stimulation.

### VLIUS promoted glymphatic function by activating TRPV4

A previous study showed that mechanical waves (shockwave or ultrasound) can activate the TRPV4 mechanosensitive channel, promoting Ca^2+^ influx into vascular endothelial cells, ultimately affecting blood-brain barrier (BBB) integrity(*25*). We wanted to know whether VLIUS, which had much lower intensities than previously used, also had similar effects on the TRPV4 channel. Therefore, we used a micropipette-guided ultrasound device(*29*) to observe the effects of VLIUS stimulation on C6 cells in vitro. Astrocytes play an important role in the regulation of glymphatic function. C6 cells are astrocyte-like cells that endogenously express TRPV4 protein. TRPV4 activation induces Ca^2+^ influx(*30, 31*). Thus, the influences of VLIUS stimulation and TRPV4 antagonists on Ca^2+^ influx were investigated to determine whether VLIUS activates TRPV4. The results showed that VLIUS stimulation promoted Ca^2+^ influx and pretreatment with TRPV4 antagonists reduced these effects in a dose-dependent manner (Fig. 3**A**).

**Fig. 3:**
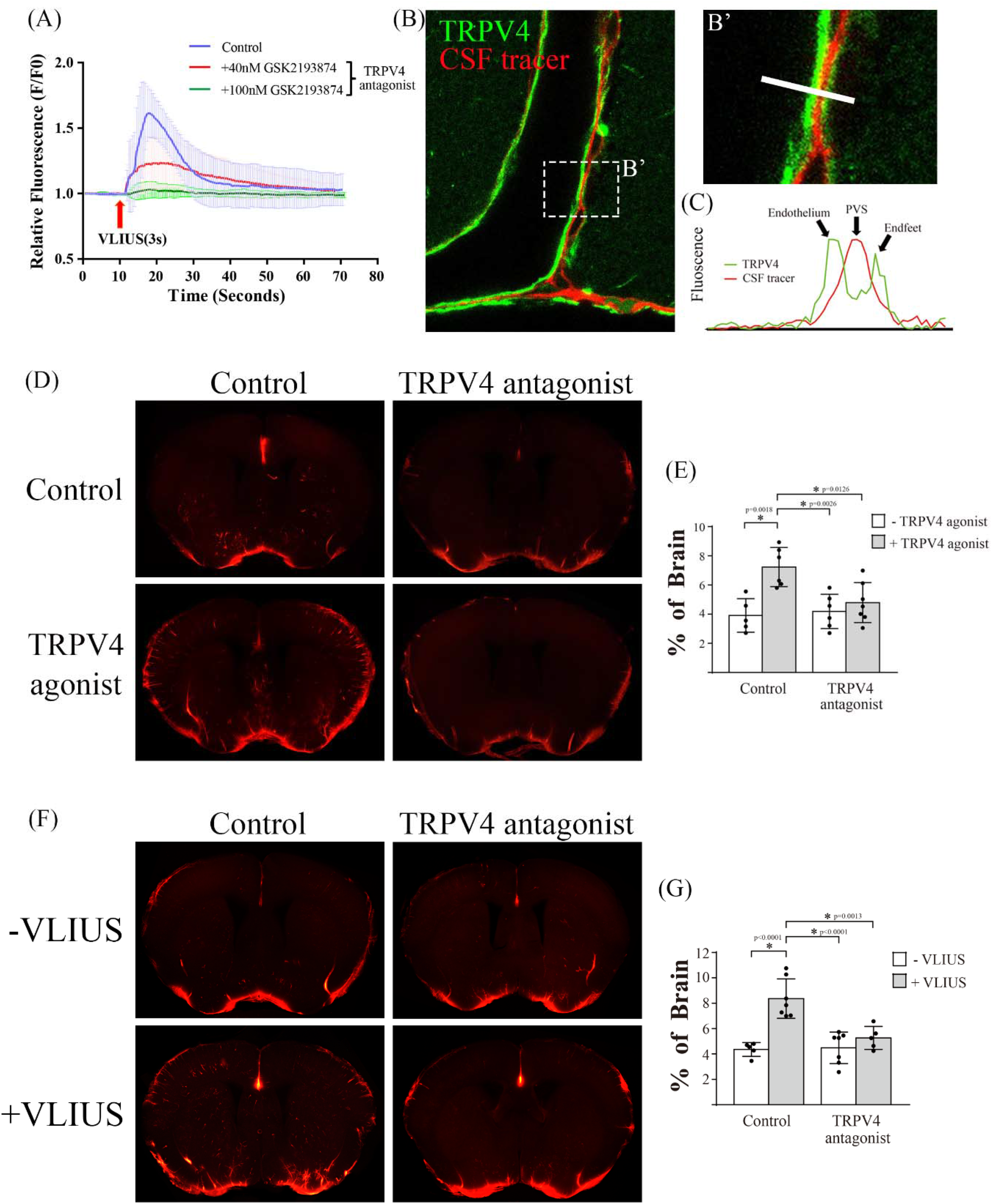
VLIUS promoted glymphatic function through activation of TRPV4. (**A**) Calcium influx elevated by VLIUS stimulation is inhibited in a dose-dependent manner by the TRPV4 antagonist. (**B** and **B’**) Along the cortical surface arteries, the CSF tracer (red) is present in the paravascular space, and TRPV4 (green) is expressed on endothelium and astrocytic endfeet. (**C**) Representative images depicting fluorescence intensity projections from (**B’**), indicated by white rectangles. (**D**) TRPV4 agonist (GSK1016790A) promotes CSF tracer permeability but is inhibited by the co-administered TRPV4 antagonist (GSK2193874). (**E**) Quantification of the influx area of a various group from (D); the dots represent individual mice in each group (control group n = 5, TRPV4 agonist group n = 6, TRPV4 antagonist group n = 6, TRPV4 agonist + antagonist group n = 7). (**F**) VLIUS-facilitated CSF permeability is blocked by TRPV4 antagonist. (**G**) Quantification of the influx area of a various group from (**F**); the dots represent individual mice in each group (control group n = 5, VLIUS group n = 7, TRPV4 antagonist group n = 7, TRPV4 VLIUS + antagonist group n = 5). Significant differences (analysis of variance with post-hoc Tukey’s test) are indicated with asterisks. VLIUS, very low intensity ultrasound; TRPV4, transient receptor potential vanilloid-4; CSF, cerebrospinal fluid.

In addition, TRPV4 is expressed at astrocytic endfeet to regulate Ca^2+^ oscillations to mediate vasodilation and vasoconstriction(*32-34*). The fact that cerebral artery pulsation drives the circulation of the glymphatic system(*35*) and that TRPV4 directly regulates vasodilation suggests that TRPV4 may affect glymphatic function. In this study, we found that the CSF tracer entered the paravascular space between the vascular endothelium and astrocyte endfeet, where TRPV4 was expressed (Fig. 3**B, B’**). Our findings are consistent with those of previous studies(*32, 36*).

We also found that tracer influx through the glymphatic system was significantly promoted by a TRPV4 agonist (GSK1016790A) and inhibited by a TRPV4 antagonist (GSK2193874) (Fig. 3**D, E**). Notably, VLIUS-stimulated glymphatic influx was inhibited by the TRPV4 antagonist (GSK2193874) (Fig. 3**F, G**), indicating the crucial role of TRPV4 in VLIUS-stimulated glymphatic influx.

### AQP4 water channel mediated TRPV4-facilitated glymphatic function

AQP4 is an important molecule that regulates the glymphatic system(*21, 37*). TRPV4 and AQP4 synergistically regulate cellular or tissue functions, such as astrocyte volume regulation(*23, 38*), calcium homeostasis(*38*), CSF secretion(*39, 40*), and make astrocytes more sensitive to extracellular osmotic gradients(*41, 42*). We hypothesized that AQP4 is involved in TRPV4-promoted glymphatic circulation. The confocal analysis of the mice brain sections revealed that TRPV4 and AQP4 co-localized at the astrocyte endfeet (Fig. 4**A, A’, B**). The TRPV4 agonist-promoted CSF tracer influx was inhibited by the concomitant administration of AQP4 inhibitors (AER271) (Fig. 4**C, D**).

**Fig. 4:**
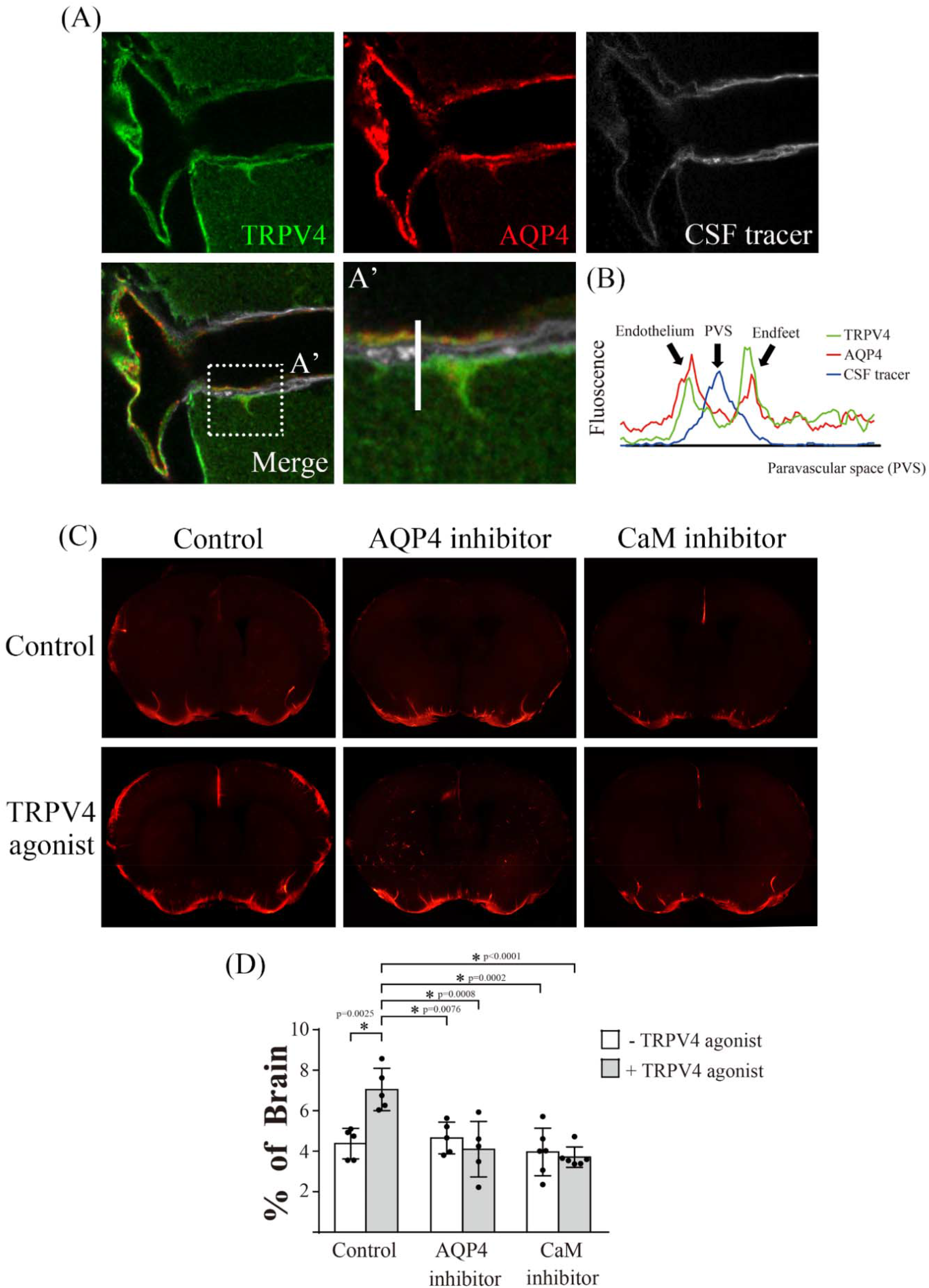
AQP4 was involved in TRPV4-facilitated glymphatic circulation. (**A**) TRPV4 and AQP4 colocalized within the paravascular space of an adult mouse brain. In single-plane confocal immunofluorescence images of cerebral surface artery, triple labeling with rabbit anti-TRPV4 (green), mouse anti-AQP4 (red), and CSF tracer (white) reveals astrocyte endfeet and endothelial cells processes that are immunopositive for TRPV4 and AQP4. (**B**) Representative images depict fluorescence intensity projections from (**A’**), indicated by the white line. (**C**) Fluorescence imaging of coronal sections reveal that 30 minutes after intracisternal injection, paravascular CSF influx has increased by administration of TRPV4 agonists and has not increased by the co-administration of AQP4 inhibitor (AER271) or calmodulin inhibitor (trifluoperazine). (**D**) Quantification of the area covered by tracer influx in coronal brain slices 30 minutes after intracisternal injection; the dots represent individual mice in each group (control group n = 5, TRPV4 agonist group n = 5, AQP4 inhibitor group n = 5, AQP4 inhibitor + TRPV4 agonist group n = 5, CaM inhibitor group n = 6, CaM inhibitor + TRPV4 agonist group n = 6). Significant differences (analysis of variance with post-hoc Tukey’s test) are indicated with asterisks. TRPV4, transient receptor potential vanilloid-4; AQP4, aquaporin-4; CSF, cerebrospinal fluid.

In contrast, the TRPV4 channel facilitates the influx of Ca^2+^ into astrocytes, thereby activating CaM, which then directly or indirectly increases the translocation of AQP4 to the cell surface to induce edema(*22, 43*). Trifluoperazine (TFP), a CaM antagonist, significantly inhibits AQP4 translocation to the cell surface in vitro and CNS edema(*22*). In this study, the TFP treatment significantly decreased the TRPV4 agonist-promoted CSF tracer influx (Fig. 4C, D). Taken together, TRPV4 agonist-promoted glymphatic function was regulated by AQP4, which was the downstream of the TRPV4-activated pathway.

### VLIUS-promoted glymphatic function regulated by AQP4 water channels

As previously described, VLIUS promoted glymphatic influx by activating TRPV4 (Fig. 3D–G), and AQP4 was required to regulate the influx of CSF tracers into the brain parenchyma after TRPV4 activation (Fig. 4C, D). We then investigated whether the VLIUS-promoted CSF tracer influx was similarly mediated by the AQP4 water channel. Concomitant administration of an AQP4 inhibitor significantly inhibited the VLIUS-induced CSF tracer influx (Fig. 5). Based on these results, the glymphatic system may be regulated by the VLIUS-TRPV4-AQP4 pathway.

**Fig. 5:**
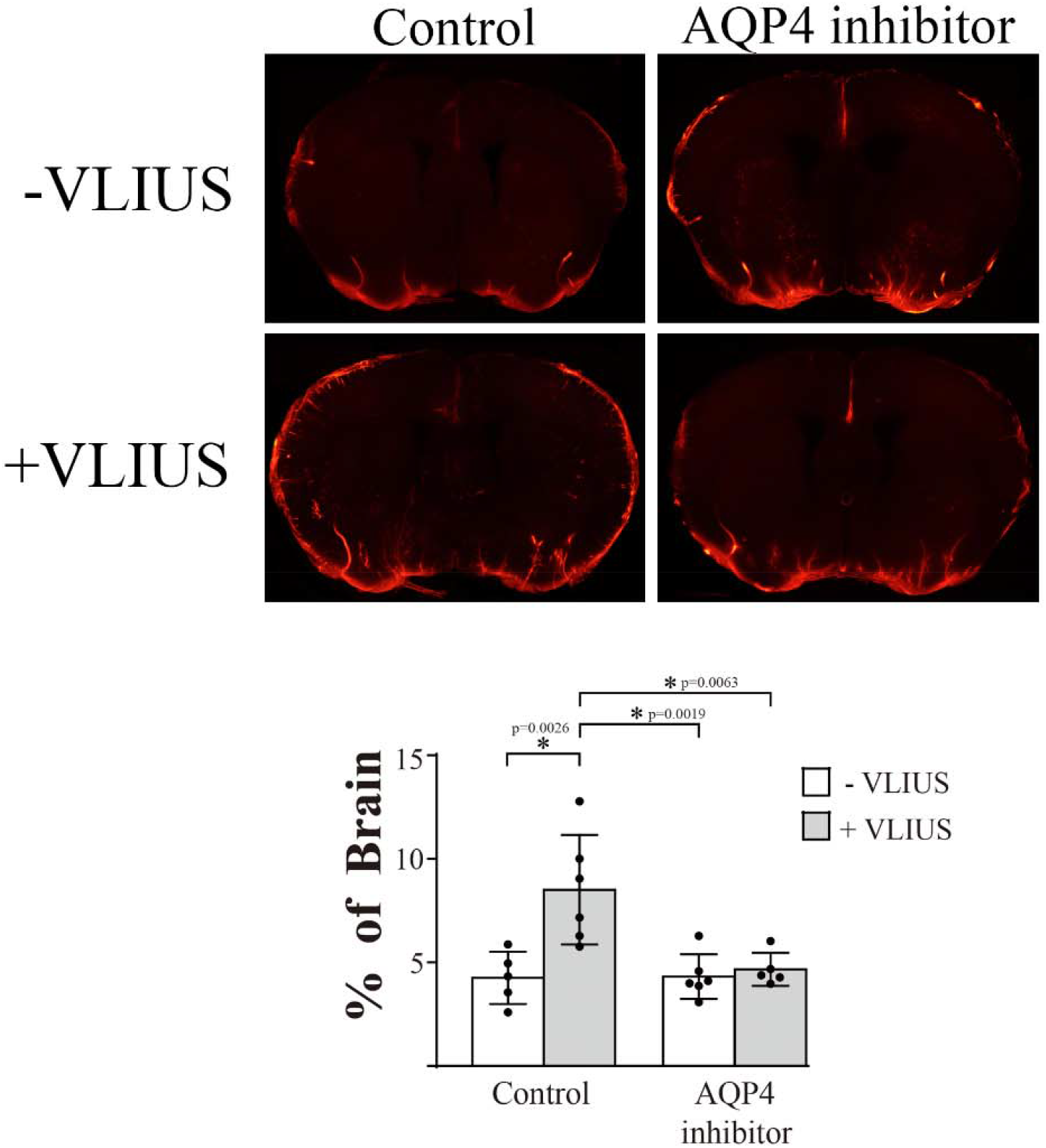
The function of AQP4 is involved in VLIUS-induced glymphatic circulation. (**A**) Fluorescence imaging of coronal sections reveals that 30 minutes after intracisternal injection, paravascular CSF influx has increased by VLIUS stimulation and has not increased by the co-administration of AQP4 inhibitor. (**B**) Quantification of the area covered by tracer influx in coronal brain slices 30 minutes after intracisternal injection; the dots represent individual mice in each group (control group n = 5, VLIUS group n = 6, AQP4 inhibitor group n = 6, AQP4 inhibitor + VLIUS group n = 5). Significant differences (analysis of variance with post-hoc Tukey’s test) are indicated with asterisks. CSF, cerebrospinal fluid; VLIUS, very low intensity ultrasound; AQP4, aquaporin-4.

### VLIUS stimulation promotes AQP4 translocation to the cell surface

Previous studies have shown that TRPV4 activation promotes Ca^2+^ influx, which promotes translocation of AQP4 to the cell surface(*22, 43*). We also demonstrated that VLIUS activated TRPV4 to promote Ca^2+^ influx into cells (Fig. 3A). Three different experiments were conducted to investigate whether VLIUS enhances the translocation of AQP4 to the cell surface through the TRPV4-mediated pathway.

First, cell surface AQP4 expression after 30-minutes of VLIUS treatment was compared to that in control cells using the cell surface biotinylation assay. The results showed that AQP4 levels at the cell surface were significantly increased by either VLIUS treatment or the TRPV4 agonist (GSK1016790A) (Fig. 6**A**). Since Ca^2+^ influx-activated CaM is a key regulator of AQP4 translocation(*22, 43*), we therefore measured the localization of AQP4 following inhibition of either CaM (using TFP) or TRPV4 (using GSK2193874). The results showed that either the CaM inhibitor or TRPV4 inhibitor decreased surface AQP4 to control levels, having reduced the effect of VLIUS on AQP4 translocation (Fig. 6A).

**Fig. 6:**
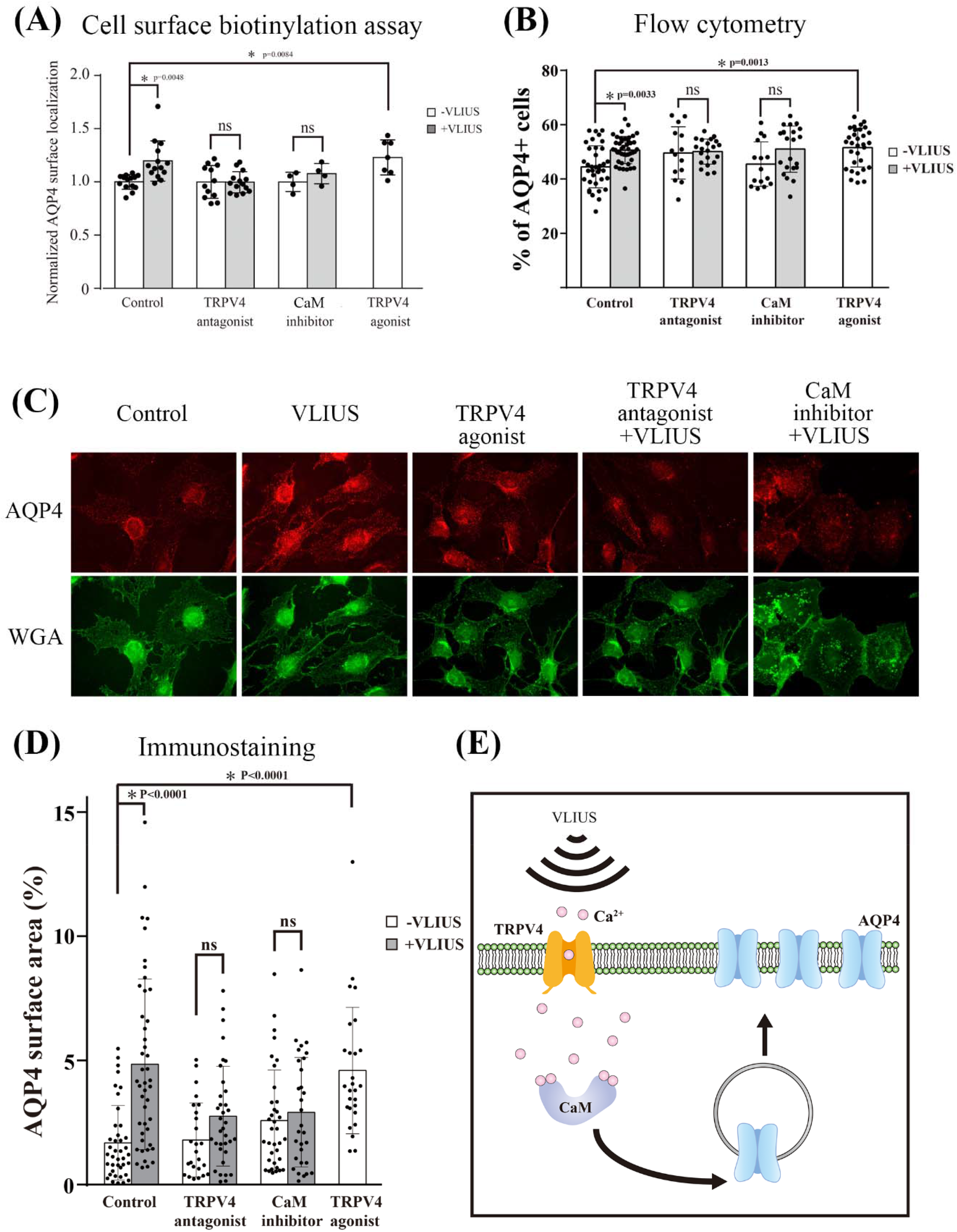
More aquaporin-4 (AQP4) protein translocated to cell surface after very low intensity ultrasound (VLIUS) stimulation. The mean fold change in AQP4 surface expression, measured by cell-surface biotinylation in C6 cells. The calmodulin (CaM) inhibitor is 20.8 μM trifluoperazine. The transient receptor potential vanilloid-4 (TRPV4) antagonist is 100 nM GSK2193874, and TRPV4 agonist is 2 μM GSK1016790A. Cells had been pre-incubated with drug 30 minutes before VLIUS treatment. AQP4 on the cell surface significantly increases after treatment with VLIUS or TRPV4 agonist. However, administration of TRPV4 antagonist or CaM inhibitor inhibit VLIUS-facilitates AQP4 cell surface localization. (**B**) The AQP4 surface expression is analyzed by flow cytometry (without permeabilization). The population of AQP4-positive cells significantly increases by VLIUS or TRPV4 agonists. Administration of TRPV4 antagonist or CaM inhibitor inhibits the VLIUS effect. (**C**) AQP4 on the cell surface is also observed by immunofluorescence staining (without permeabilization). Cells had been counterstained by WGA-conjugated Alexa Fluor™ 488 to determine cell boundaries. (**D**) Quantification of AQP4 on the cell surface using immunofluorescence staining. (**E**) VLIUS stimulation activates the mechanosensitive TRPV4 channel, then facilitates an influx of calcium ions into astrocytes, activating CaM. Significant differences (analysis of variance with post-hoc Tukey’s test) are depicted with asterisks.

Second, we used flow cytometry to analyze the proportion of AQP4^+^ cell population on the surface. The cells were only fixed with 4% paraformaldehyde (PFA) without TritonX-100 treatment before immunostaining to ensure that only the AQP4 proteins on the cell surface were detected. Both the VLIUS stimulation and TRPV4 agonist significantly increased the AQP4^+^ cell population. In contrast, both the TRPV4 antagonists and CaM inhibitors reduced VLIUS-induced AQP4 translocation (Fig. 6**B**).

Third, we observed AQP4 on the cell surface by fluorescence microscopy in the absence of TritonX-100 treatment and used wheat germ agglutinin (WGA) staining to determine the boundary of the cells. The intensity and proportion of AQP4 on the cell surface significantly increased after VLIUS treatment or TRPV4 agonist treatment (Fig. 6**C, D**). Combining the above three experiments, AQP4 protein levels on the cell surface were significantly increased after VLIUS treatment.

### VLIUS stimulation modulated astrocytic cell volume within glia limitans

We found that VLIUS stimulation promoted AQP4 translocation to the cell surface. We wanted to determine how this affected the cells. AQP4 mediates water influx to induce cell swelling and then triggers a regulatory volume decrease (RVD) to restore its original volume in response to swelling, suggesting that AQP4 is involved in regulating cell volume(*44, 45*). Furthermore, AQP4 abundance on the cell surface induces cytotoxic edema, whereas attenuation of edema through AQP4 inhibition improves electrophysiological, sensory, and locomotor function in animals(*22*). Since AQP4 translocated to the cell surface after VLIUS stimulation, we investigated whether VLIUS induced cell volume changes and led to persistent cytotoxic edema.

In this study, we observed glial fibrillary acidic protein (GFAP)-positive astrocytes in the glia limitans (Fig. 7**A, B**). The glia limitans was selected rather than the perivascular astrocytic endfeet because (1) the spatial structure of the endfeet is not easy to observe, distinguish, and analyze; (2) the astrocytic endfeet around the blood vessels are extensions of the glia limitans; and (3) we observed that the CSF tracer diffuses not only from the paravascular space, but also from the brain surface to the brain parenchyma with time (Supplementary Fig. S3). Furthermore, it has been reported that glia limitans are composed of astrocytes with high GFAP expression(*46*).

**Fig. 7:**
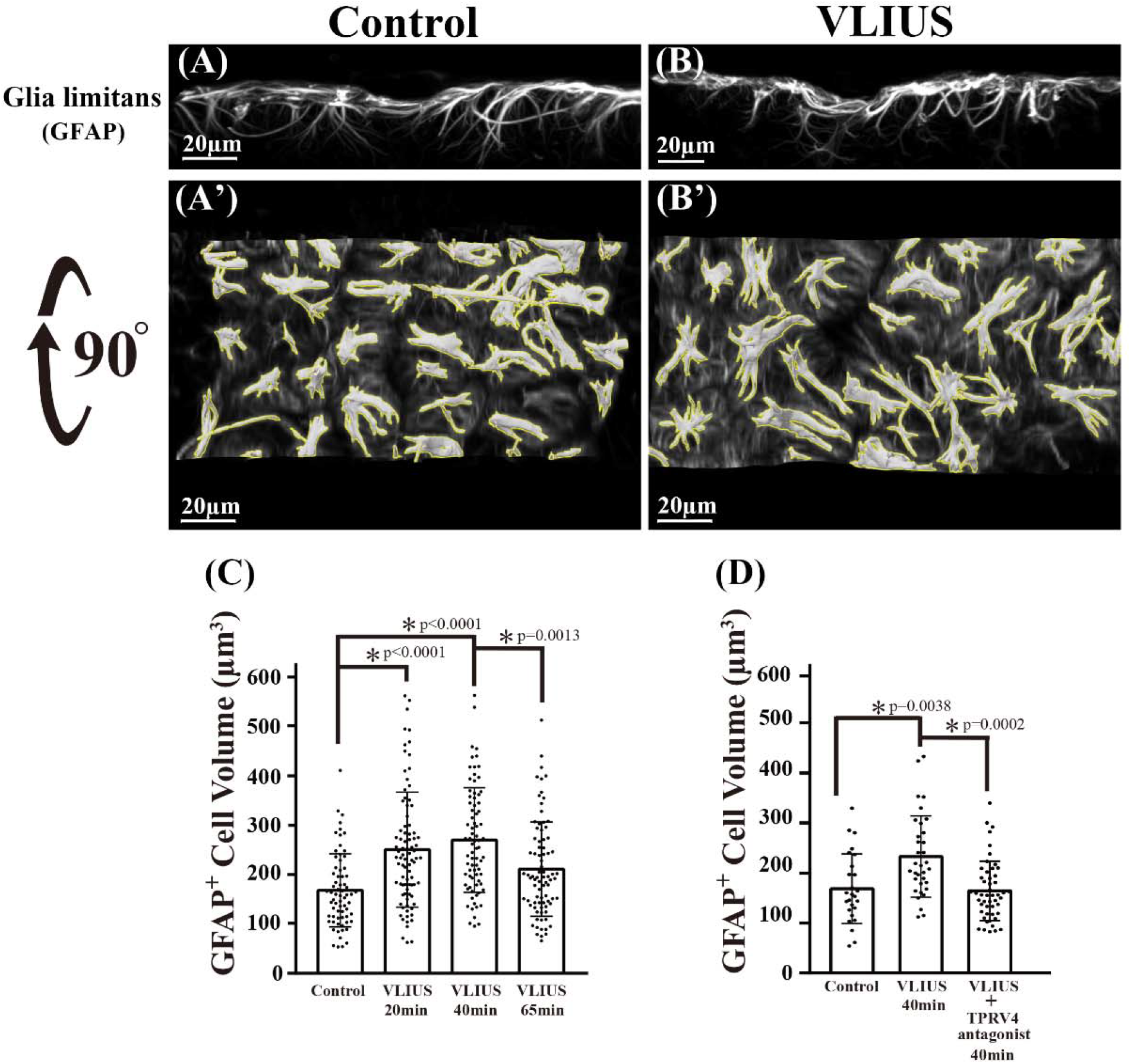
Volume of astrocytes in the glial limitans altered after very low intensity ultrasound (VLIUS) stimulation. 3D images from confocal microscopy had been reconstructed using Imaris software, and are shown as control (**A**), VLIUS (**B**), and 90° rotated images (**A’**) and (**B’**). The yellow line represents the analysis of each intact cell (with the same threshold) using the surface creation wizard of the Imaris software. Quantification shows that cell volume is altered by VLIUS stimulation (**C**) and the effect is blocked by the transient receptor potential vanilloid-4 (TRPV4) antagonist (**D**). The dots represent individual intact cells in each group. Significant differences (analysis of variance with post-hoc Tukey’s test) are indicated with asterisks.

We attempted to understand the role of AQP4 translocation to the cell surface by observing changes in the volume of GFAP-positive astrocytes. The volume of GFAP-positive cells significantly increased at 20 and 40 minutes after VLIUS stimulation and decreased significantly at 65 minutes after VLIUS stimulation (Fig. 7**A**–**C**). In other words, the volume of GFAP-positive cells increased transiently (approximately 20–40 minutes) after VLIUS stimulation and was restored later, implying that VLIUS stimulation did not induce persistent cytotoxic edema. Furthermore, we did not observe any harmful effects after VLIUS stimulation (Supplementary Fig. S4).

Both TRPV4 alone(*47*) and TRPV4 with AQP4(*23, 38*) modulate cell volume and RVD. The expression of TRPV4 and GFAP co-localized within glial limitans(*48*). In the current study, we found that TRPV4 activation enhanced AQP4 translocation to the cell surface (Fig. 6), and that the TRPV4 antagonist significantly inhibited VLIUS-induced increases in cell volume (Fig. 7**D**). Therefore, TRPV4 may be activated by VLIUS stimulation to increase the translocation of AQP4 to the cell surface and thereby modulate water into and out of the cell to alter cell volumes.

## DISCUSSION

The major finding of this study was that VLIUS enhanced glymphatic influx via the TRPV4-CaM-AQP4 pathway in astrocytes. TRPV4 was activated by VLIUS in the astrocytes to induce a Ca^2+^ influx to activate CaM, which promoted the translocation of AQP4 to the cell surface, inducing water influx, and thus increased cell volume. Sixty-five minutes after VLIUS stimulation, the swollen cells were restored to their original volume, possibly due to the efflux of water. This transcellular water flow may be the driving force behind VLIUS-stimulated glymphatic circulation.

The glymphatic system is considered an important target for therapeutic intervention in CNS disorders(*27, 49*). Ultrasound, due to its non-invasiveness and availability, has been applied for the treatment of various CNS disorders, with the outcome effects mainly focusing on opening the BBB. However, the incident ultrasound intensity or pressure is usually much higher than that used in this study. Furthermore, most studies have added microbubbles (MBs) to generate cavitation and open the BBB. Thus, brain injury risk is increased. If the generated cavitation is too strong or if a patient’s blood vessels or BBB structures are too fragile, irreversible damage may occur(*50*). In the current study, a planar ultrasound transducer was used and the intensity was relatively low (I_spta_: 3.68 mW/cm^2^, pressure level 36 kPa). As previously shown, low energy can still generate various bioeffects, such as promoting adult neurogenesis in the dentate gyrus(*28*). Another study focusing on brain neurogenesis in the dentate gyrus also showed that the group without MBs had better bioeffects and functional recovery than the group with MBs(*51*), implying that MBs are not necessary for brain stimulation. In the current study, we showed that VLIUS without MBs enhanced glymphatic influx and no harmful effects on the brain were observed (Supplementary Fig. S4). VLIUS, without MBs, can be a safe and effective way to modulate the glymphatic system.

The TRPV4-CaM-AQP4 pathway may be one of the mechanisms through which VLIUS enhances glymphatic function. One study indicated TRPV4 interacts with AQP4 to regulate the entry and exit of water molecules in and out of cells for regulating cell volume, which promotes Ca^2+^ influx through TRPV4 activation and subsequently regulates RVD in hypotonic environments(*23*). However, another study suggested TRPV4-induced Ca^2+^ influx was not important because TRPV4 blockade or removal of external Ca^2+^ did not prevent RVD in hypotonic environments(*45*). The authors used an overexpression of AQP4-M23 to observe cell volume changes, allowing water to enter cells spontaneously(*45*). Since AQP4 has multiple isoforms that are present in cells in different proportions and play different roles(*44*), overexpression of AQP4-M23 alone may not explain the entire condition. Furthermore, the response generated by residual Ca^2+^ cannot be completely excluded(*45*). Previous studies have suggested that TRPV4 induces Ca^2+^ oscillation in astrocyte endfeet to regulate neurovascular coupling(*32-34*). Activated TRPV4 promotes external Ca^2+^ influx into cells and triggers the activation of inositol trisphosphate receptors (IP3Rs), amplifying local Ca^2+^ signals and initiating Ca^2+^ waves(*23*). Only a small Ca^2+^ influx is sufficient to amplify Ca^2+^ signals and form subsequent Ca^2+^ oscillations. Therefore, it is generally considered that the synergistic effect of TRPV4 and AQP4 regulates cell volume and increases the sensitivity of astrocytes to environmental stress(*22, 41*). However, these studies were based on hypotonicity stress, which activates TRPV4 due to changes in cell membrane tension caused by AQP4, facilitating water entry into cells. In our study, the ultrasound-generated mechanical force to activate TRPV4 may have differed from hypotonic-induced changes in membrane tension.

Activation of TRPV4 by agonists or VLIUS can promote the transport of AQP4 from the cytosol to the cell surface (Fig. 6)(*22, 43*) which can effectively promote CSF tracer influx, while TRPV4 blockade significantly inhibited VLIUS-promoted CSF tracer influx (Fig. 3). Our results also showed that the effects of VLIUS stimulation were better than those of TRPV4 agonists alone, which suggests that besides TRPV4 receptors, other mechanosensitive channels sensitive to VLIUS stimulation may be present in astrocytes(*52*). Moreover, prolonged activation of TRPV4 by agonists can lead to irreversible side effects such as reduced expression levels of tight junction proteins(*53, 54*), BBB disruption(*54*), and apoptosis induction(*55, 56*). Unlike CNS injury, VLIUS stimulation did not cause irreversible cytotoxic edema. The increased cell volume stimulated by VLIUS returned to baseline 65 minutes later (Fig. 7). No tissue disruption was detected in the tissue staining assays (Supplementary Fig. S4). Therefore, VLIUS may be a better choice for promoting glymphatic circulation.

This study has several clinical implications. First, we observed that VLIUS, without MBs, enhanced glymphatic function without imposing brain injuries in mice. Since the brain volume and skull thickness are different in humans and mice, modification of certain VLIUS parameters is necessary for VLIUS to provide positive effects in human CNS disorders. Second, the glymphatic system has a low function on awakening. VLIUS may be incorporated into a wearable device and provide a continuous enhancement of glymphatic clearance even on awakening. Before this, any harmful effects of all day long glymphatic enhancement should be excluded.

This study has several limitations. First, AQP4 agonists were not available; therefore, we were unable to confirm whether activation of AQP4 affects TRPV4 or not. Which one is upper stream, AQP4 or TRPV4, remains controversial. Second, in postmortem samples the arteries collapse, and the fluid in the perivascular space largely disappears(*37*), which could lead to misjudgment in observation. Using transgenic mice with fluorescent astrocytes under 2-photon microscopy observation would help visualization of more details(*57*).

The exact mechanism of RVD modulation cannot be explained based on our results. It is speculated that the driving force generated by Cl^-^ efflux via activation of downstream volume-regulated anion channels or calcium-activated chloride channels causes water efflux and reduces cell volume(*39, 58*). TRPV4, AQP4, and anoctamin 1 cooperate to regulate water efflux and cell volume of choroid plexus epithelial cells (CPECs), suggesting their roles in CSF production(*39*). Furthermore, TRPV4 appears to be a cell surface sensor that promotes cell secretion after receiving extracellular stimuli. Activation of TRPV4 promotes the exocytosis of various molecules, including AQP4 (in astrocytes)(*22*), TRPV4 itself (in human umbilical vein endothelial cells)(*59*), alpha-klotho, and sodium, potassium-adenosine triphosphatase (in CPECs)(*60*). Notably, upon stimulation, TRPV4 is activated to promote the secretion of water or proteins from CPECs into the extracellular space, implying that TRPV4 may regulate CSF secretion. Since TRPV4 is abundantly expressed in the choroid plexus, VLIUS stimulation may also promote CSF secretion. Therefore, further studies are warranted.

## MATERIALS AND METHODS

### Animals

All animal experiments were carried out in accordance with the Institutional Animal Care and Use Committee guidelines of the National Taiwan University College of Medicine (IACUC approval No. 20201028). C57BL/6JNarl mice (body weight, 20–25 g) purchased from the National Laboratory Animal Center (Taiwan) were used in this study. The mice were maintained in a controlled 12-hour light/12-hour dark cycle with access to water and food ad libitum. In all experiments, animals were anesthetized with a combination of zoletil (50 mg/kg, intraperitoneally) and xylazine (2.3 mg/kg, intraperitoneally)

### Ultrasound devices and stimulation parameters

The ultrasound device setup has been described in our previous study(*61*). The ultrasound field was generated using a 1 MHz commercial plane transducer (C539-SM, Olympus, Tokyo, Japan) for mouse brain stimulation. All stimulations were performed using a function generator (Tektronix AFG1022, Beaverton, OR, USA) through a power amplifier (E&I 210 L, Electronics & Innovation, Rochester, USA). The planar transducer was placed at the center of the brain, and an ultrasound gel was applied at the interfaces between the bottom of the transducer and the mouse scalp. After intracisternal injection, mice were immediately stimulated with VLIUS for 5 min. VLIUS treatment was performed under the following conditions: center frequency = 1 MHz, pulse repetition frequency = 1 kHz, duty factor = 1 %, and various spatial peak temporal average intensities (I_spta_) = 0.92, 3.68, and 5.85 mW/cm^2^ (Supplementary Fig. S1, Fig. S2).

Among them, I_spta_ = 3.68 mW/cm^2^ (pressure level = 36 kPa) was the best condition for promoting tracer diffusion (Supplementary Fig. S2); therefore, subsequent experiments were all stimulated under this condition.

For in vitro experiment, cells were seeded 1 × 10^5^ cells on a 24 well plate 18 hours before the experiments. After the medium was renewed, a planar transducer was applied directly above the cells in the medium. The VLIUS setting was the same as described above, except that the duration was changed to 1 minute. One hour after VLIUS stimulation, the cells were washed with phosphate buffered saline (PBS) and subsequent experiments were performed. For micro-pipette-guided ultrasound stimulation, please refer to the section on live-cell calcium signal imaging.

### Intracisternal CSF tracer infusion

Intracisternal injection was performed as previously described with a minor modification(*62*). Mice were weighed and anesthetized by intraperitoneal injection of zoletil/xylazine. The cisterna magna was exposed through a surgical incision using a stereotactic frame, and a 30-gauge needle connected to a Hamilton syringe via a polyethylene tube (PE10) was inserted into the cisterna magna. Fluorescent CSF tracer (albumin-Alexa Fluor™ 555, Cat No. A34786, Thermo Fisher Scientific, Waltham, MA, USA) was added to the artificial CSF at a concentration of 5 mg/ml. A CSF tracer (6 µl) was infused at a constant rate of 1 µl/minute with a syringe pump (Harvard Apparatus, Holliston, MA, USA). During the experiment, the needle was kept in place to avoid depressurization of the CSF compartment. In the VLIUS stimulation group, the mice were immediately stimulated with VLIUS for 5 minutes after intracisternal infusion. Thirty minutes after the start of intracisternal infusion, the mice were euthanized, and brain was harvested and fixed with 4% cold paraformaldehyde in PBS overnight. For evaluating the distribution of tracers into the brain parenchyma, the 500 µm coronal slices were cut on a vibratome (MicroSlicer™ DTK-1000N, DSK, Kyoto, Japan) and then incubated in 2% phosphate-buffered saline with Tween (PBST) (2% Triton X-100 in PBS) for 2 days to promote tissue permeability. After washing three times with PBS, the tissues were cleared and incubated with RapiClear ® 1.49 (SunJin Lab Co., Hsinchu City, Taiwan) for 1 hour to promote transparency and then mounted with fresh RapiClear®. Samples were imaged using a fluorescence microscope (Olympus BX51 with ToupTek camera) and the extended depth of field method (ToupView software, Hangzhou, China) to provide higher-quality images and faster acquisition of large amounts of information. Tracer influx (area %) was analyzed using ImageJ software (16-bit image type; analyze particle setting: 80–infinity µm^2^ size range, 0.0–1.0 circularity, and automatic threshold). The number of paravascular influxes was counted in the brain section where the subarachnoid tracer had extended into the parenchyma (Fig. 1C). The influx length was measured from the subarachnoid space to the distal end using ToupView software (Fig. 1D). Additionally, to measure the diffusion of the tracer in the anterior to posterior regions of the brain, we used 100 µm brain slices instead for more refined results (Fig. 1E).

### Transcranial live imaging

For in vivo transcranial live imaging, the skin covering the dorsal calvarium was incised and a tracer was injected through the cisterna magna. The entry of CSF tracers into the brain was imaged using fluorescence macroscopy (Cat No. MVX10, Tokyo, Olympus) with an ORCA-Spark digital CMOS camera (Cat No. C11440-36U, Hamamatsu, Japan). Images were recorded at 30 second intervals for 0–64 minutes following injection commencement using the CellSens Standard 3 (Olympus). The exposure time was the same throughout the imaging sequence and across the experimental groups.

### Intraparenchymal tracer injection

To evaluate the clearance rates of interstitial fluid from the brain after VLIUS treatment, the tracer was stereotactically injected into the brain parenchyma. The anesthetized mice were fixed in a stereotaxic frame, and a 32G needle was inserted via a burr hole into the left striatum (0.22 mm caudal, 2.5 mm lateral, 3.5 mm ventral to bregma). The tracer (total 1 µl) was injected at a rate of 200 nl/minute, and the needle was left in place for an additional 30 minutes. After intrastriatal injection, the mice were treated with or without VLIUS stimulation, then a 3 hour wait time occurred to allow the tracer to be cleared from the parenchyma through the interstitial solute clearance mechanism. After 3 hours, the mice were euthanized, and the brains were harvested. To measure the tracer content, brain tissues were homogenized on ice with the Pierce™ RIPA Buffer (Cat No. 89900, Thermo Fisher Scientific), then centrifuged at 12,000 g at 4 °C for 20 minutes. The supernatants were selected and measured using a microplate reader (Infinite M200; Tecan, Austria). The tracer content is shown in relative light units per milligram of protein. The protein concentration was measured using the Pierce™ Coomassie Protein Assay (Cat No. 1856209, Thermo Fisher).

### Drug administration

#### In vivo study

The related drugs, doses, timing, and modes of administration are shown in Table 1. Before intracisternal infusion, the experimental animals received a single injection of drugs dissolved in 0.1 ml normal saline. The control animals received 0.1 ml of normal saline.

#### In vitro study

The related drugs, doses, and timing are shown in Table 2. The cells were pre-incubated with the drugs for 30 minutes before VLIUS treatment.

### Free-floating immunofluorescence staining

For free-floating immunofluorescence staining, the 250 µm coronal slices were cut on a vibratome (MicroSlicer™ DTK-1000N, DSK) and then incubated in 2% PBST (2% Triton X-100 in PBS) for 1 day to promote tissue permeability. Slices were washed three times with PBS and blocked with fresh blocking buffer (10% normal goat serum, 1% Triton-X100, 2.5% DMSO, and 0.2% sodium azide) for 1 day on a rocker at 4 LJ. The slices were incubated with primary antibody in SignalStain^®^ antibody diluent (Cell Signaling, Cat No. 8112, USA) on a rocker for 2 days at 4 LJ, and then washed three times with washing buffer (3% NaCl and 0.2% Triton-X100 in PBS) for 1 hour at room temperature (RT). The slices were kept in washing buffer on a rocker at 4 LJ overnight. The slices were incubated with secondary antibody on a rocker for 2 days at 4 LJ, and then washed three times with washing buffer, and kept in washing buffer on a rocker at 4 LJ. After washing with PBS (three times), clear samples with RapiClear ® 1.49 (SunJin Lab Co., Hsinchu City, Taiwan) for 1 hour to promote transparency and then mounted with fresh RapiClear ®. The primary antibodies used were rabbit anti-TRPV4 (Cat No. PA5-41066, 1:100, Thermo Fisher Scientific), mouse anti-GFAP (Cat No. MA5-12023, 1:500, Thermo Fisher Scientific) and mouse anti-AQP4 (Cat No. sc-32739, 1:100, Santa Cruz Biotechnology, Dallas TX, USA). The secondary antibodies used were Alexa Fluor 633-conjugated goat anti-mouse antibody (Cat No. A-21050, 1:200, Thermo Fisher Scientific) and Alexa Fluor 488-conjugated goat anti-rabbit antibody (Cat No. A-27034, 1:200, Thermo Fisher Scientific). Samples were imaged using laser scanning confocal microscopy (Carl Zeiss LSM880) and analyzed using ImageJ software.

To analyze astrocyte volume within glia limitans, GFAP expression was further observed by laser scanning confocal microscopy at a 63X objective to record fluorescence images of the glia limitans. These were acquired in 70–100 µm z-stacks (∼0.69 µm z-steps). Cell volume was evaluated and analyzed using the Imaris software (Oxford Instruments, 9.5 version, UK). The Z-stack images were opened in Imaris in their native format. Z-stacks were automatically reconstructed into a 3D model using Imaris software without image preprocessing. The surface creation wizard was used to analyze the volume of each intact cell (with the same threshold).

### Cell culture

C6 glioma cells (Bioresource Collection and Research Center, Hsinchu City, Taiwan), an immortalized rat glial cell line, were maintained in Dulbecco’s modified Eagle’s medium containing high glucose (DMEM) (Cat No. 12100046, Thermo Fisher Scientific) supplemented with 10% fetal bovine serum (Thermo Fisher Scientific) and a 1 × antibiotic–antimycotic (Cat No. 15240112, Thermo Fisher Scientific) at 37 °C in 5% CO_2_. Western blot analysis showed that C6 cells natively expressed TRPV4 and AQP4 (data not shown). In addition, C6 cells are considered to have astrocytic properties(*63*). Therefore, C6 cells were selected for the subsequent in vitro studies.

### Live cell calcium signal imaging

For the calcium influx assay, we used a micro-pipette-guided ultrasound system(*29*). Cells (3 × 10^5^) were seeded on a 30 mm circular coverslip 18 hours before the experiments. Cells were washed twice with Hank’s Balanced Salt Solution (HBSS) (Cat No. 14025134, Thermo Fisher Scientific) and incubated with 5 µM Fluo-8 AM (Cat No. ab142773, Abcam, Bristol, UK) 30 minutes. Afterwards, the cells were washed three times with HBSS and incubated in serum-free DMEM (no phenol red, Cat No. 21063029, Thermo Fisher Scientific) for 30 minutes at RT. The coverslip was mounted on the microscopy chamber and placed under an Olympus IX71 microscope, and a micropipette ultrasound was set up near the targeted cells. Calcium influx images were recorded with the same exposure protocol: 10 seconds pretreatment followed by a 3 second ultrasound stimulation and then a 57 second wait time. Stack images were analyzed with ImageJ software. Areas of interest were determined for the stacks, and a graph of the fluorescence intensity against time points was plotted.

### Cell surface biotinylation

C6 glioma cells were plated in 24 well plates 1 d before each experiment. Cell surface proteins were biotinylated using the EZ-Link™ Sulfo-NHS-SS-Biotin reagent (Cat No. 21331, Thermo Fisher Scientific). Cells were treated in three experimental conditions (VLIUS, agonist, antagonist, or inhibitor) and then incubated in 250 µL of 0.5 mg/ml biotinylation reagent in PBS on ice for 30 minutes. The unlabeled reagent was quenched with 25 mM glycine in PBS per well for 3 × 5 minutes. Cells were lysed in 100 µL Pierce™ RIPA Buffer (Cat No. 89900, Thermo Fisher Scientific) supplemented with protease inhibitor cocktail set III (Cat No. 539134-1 set, 1:100, Merck Millipore, Rahway, NJ). The lysate was centrifuged at 16,000 g at 4 LJ for 10 minutes to remove insoluble samples. Each lysate was measured using the Coomassie protein assay reagent (Cat No. 1856209, Thermo Fisher Scientific) for normalization. Biotinylated proteins were immobilized out by incubation in 96-well high-sensitivity streptavidin microplates (Cat No. 6523-5, BioVision, Waltham, MA, USA) for 2 hours at 4 LJ with shaking. Plates were blocked with 3% w/v bovine serum albumin (BSA) in PBS for 1 hour at RT with shaking and then incubated on a shaker overnight at 4 LJ with the anti-AQP4 antibody (Cat No. GTX133151, GeneTex, San Antonio, TX, USA) diluted 1:400 in 0.1% PBST 20. Plates were washed three times with 0.1% PBST and incubated at RT for 1 hour with horseradish peroxidase-conjugated secondary antibody (Cat No. NA934, Merck Millipore) diluted 1:1,000 in 0.05% PBST. Plates were washed with 0.1% PBST five times then once with PBS, and incubated with 3,3’,5,5’-tetramethylbenzidine substrate solution (Cat No. N301, Thermo Fisher Scientific) for 20 minutes (protected from light). The absorbance was measured at 450 nm using a microplate reader (Infinite M200).

### AQP4 surface membrane localization by flow cytometry analysis

One hour after VLIUS stimulation, the C6 cells were washed with cold PBS and resuspended in Accutase® cell detachment solution (Cat No. AT104, Innovative Cell Technologies, San Diego, CA, USA). Cells were fixed with 4% PFA for 15 minutes at RT, and then washed three times with PBS. Because we only detected AQP4 protein on the cell surface, the cells were not permeabilized (such as with Triton X-100 or methanol treatment), as it facilitated antibody entry into cells to detect intracellular AQP4 protein. Cells were blocked with block solution (1% w/v BSA in PBS) for 1 hour at RT and then incubated with the anti-AQP4 antibody (GeneTex, Cat No. GTX133151, diluted 1:200 in block solution) overnight at 4 LJ. Cells were washed three times with the block solution and incubated at RT for 2 hours with a secondary antibody (Cat No. A-21428, Thermo Fisher Scientific) was diluted 1:400 in a block solution. Cells were washed three times with PBS, and then analyzed using LSRII flow cytometry (BD Biosciences, East Rutherford, NJ, USA) through the service provided by the Flow Cytometric Analyzing and Sorting Core Facility at the National Taiwan University Hospital.

### AQP4 surface membrane localization by immunofluorescence staining

Cells were seeded at 5 × 10^4^ cells on a 15 mm circular coverslip (in 24 well plate) 18 hours before the experiments. After 1 hour of VLIUS stimulation, C6 cells were washed with cold PBS and fixed by incubation with 4% PFA for 15 minutes at RT. Likewise, cells were not permeabilized, and only cell surface AQP4s were detected using the antibody. Cells were blocked with 1% w/v BSA in 0.1% PBS-Tween 20 for 1 hour at RT and then incubated with an anti-AQP4 antibody (Cat No. GTX133151, GeneTex) diluted 1:400 in SignalStain^®^ antibody diluent (Cat No. 8112, Cell Signaling Technology, Danvers, MA, USA) overnight at 4 LJ. Cells were washed three times with the block solution and incubated at RT for 2 hours with a secondary antibody (Thermo Fisher Scientific, Cat No. A-21428; 1:200 in SignalStain^®^ antibody diluent). Cells were counterstained with WGA and Alexa Fluor™ 488 conjugate (Cat No. W11261, Thermo Fisher Scientific) to determine the cell range. Finally, the cells were mounted using EverBrite mounting medium with 4′,6-diamidino-2-phenylindole (Cat No. 23002, Biotium, Fremont, CA, USA). Images were recorded by Olympus IX51 microscope with ToupTek camera and with the same exposure parameters (Alexa Fluor™ 555: exposure time = 30 ms, gain = 200%; WGA-Alexa Fluor™ 488: exposure time = 10 ms, gain = 200%). Images were analyzed using ImageJ software.

### Statistics

All numerical data are expressed as mean ± standard deviation. Tests of significance were performed using analysis of variance, followed by Tukey’s post-hoc test. For all studies, p values < 0.05 were considered statistically significant. All statistical comparisons were performed using GraphPad Prism software (GraphPad Software, La Jolla, CA, USA).

## Acknowledgments

We thank the imaging core at the First Core Labs, National Taiwan University College of Medicine, for the technical support in image acquisition and analysis. We would like to acknowledge the service provided by the Flow Cytometric Analyzing and Sorting Core Facility at National Taiwan University Hospital.

## Author contributions

Conceptualization: WHL, CHW, JLW, WSC

Methodology: WHL, CHW, YCC, MYH, YK, JLW, WSC

Investigation: WHL, CHW, JLW, WSC

Visualization: WHL, CHW, YCC, MYH, YK, JLW, WSC

Funding acquisition: CHW, MYH, JLW, WSC

Project administration: WHL, CHW, YCC, MYH, YK

Supervision: JLW, WSC

Writing – original draft: WHL, CHW

Writing – review & editing: WHL, CHW, YCC, MYH, YK, JLW, WSC

## Data and materials availability

All data are available in the main text or the supplementary materials.

## Supplementary Materials

**Fig. S1:**
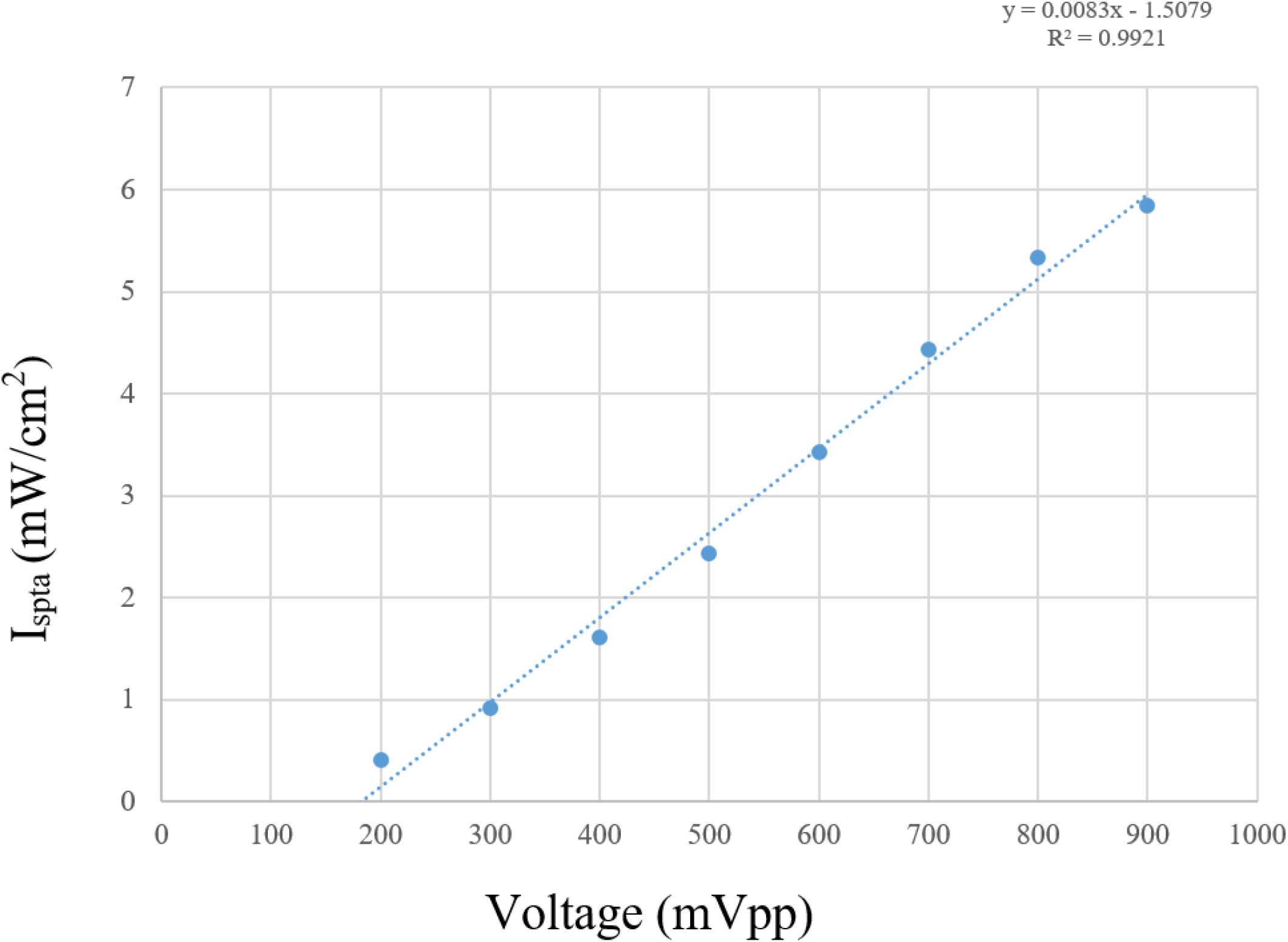
Linear correlation between spatial-peak temporal average (Ispta) and input voltage (mVpp).

**Fig. S2:**
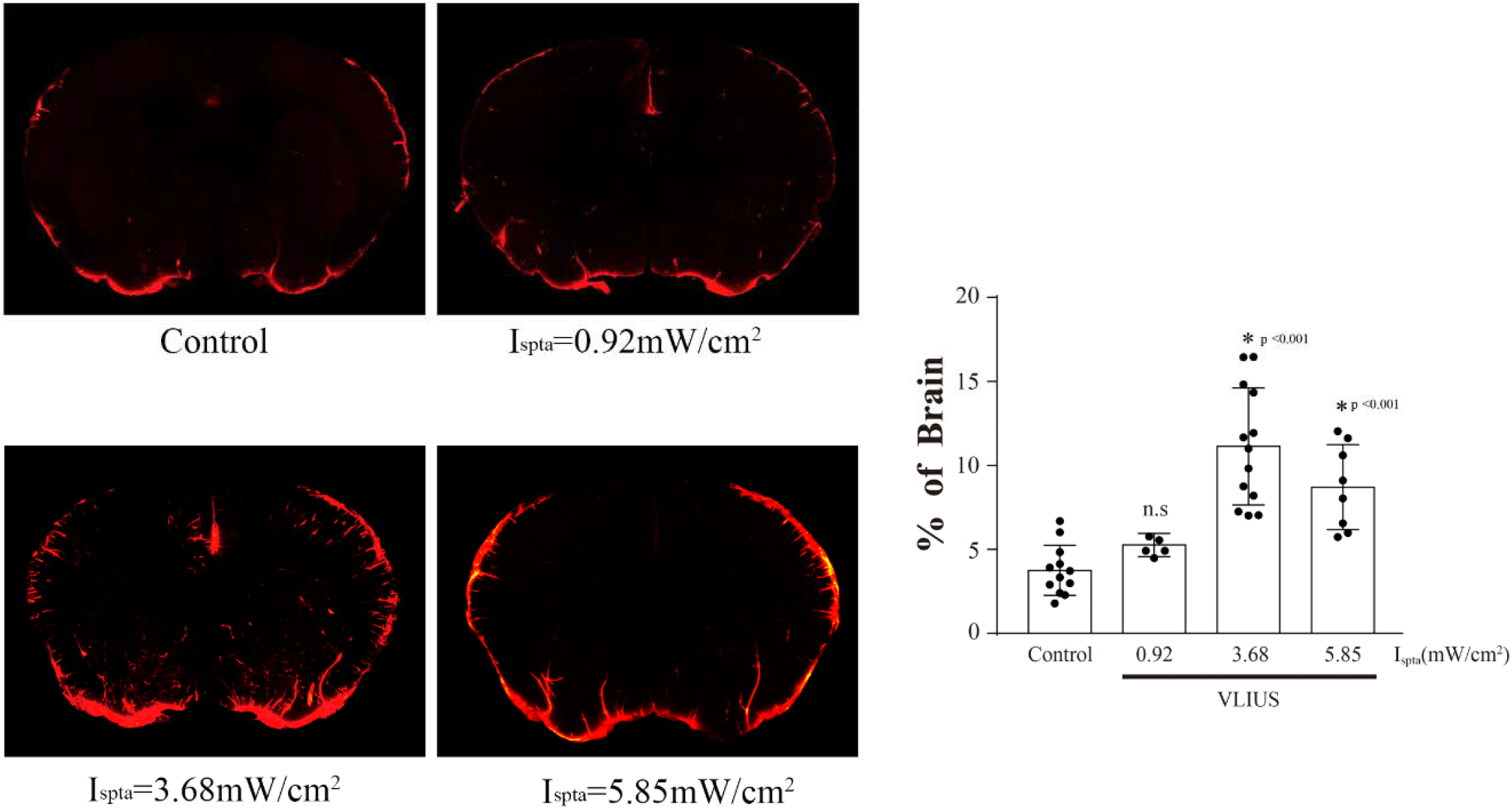
Enhancement of glymphatic system function using ultrasound at different intensities.

**Fig. S3:**
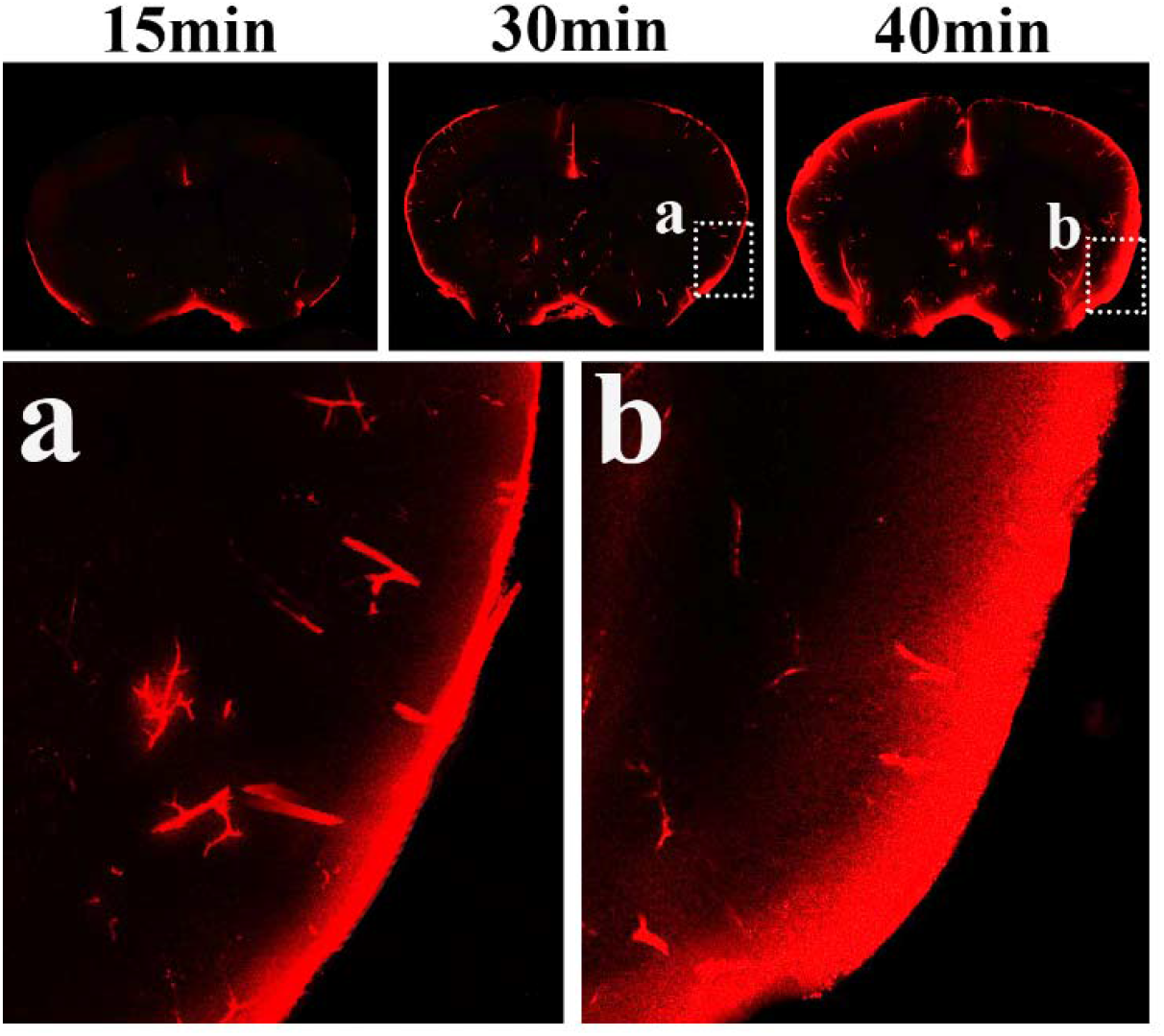
Diffusion of the tracer from the brain surface to the brain parenchyma with time.

**Fig. S4:**
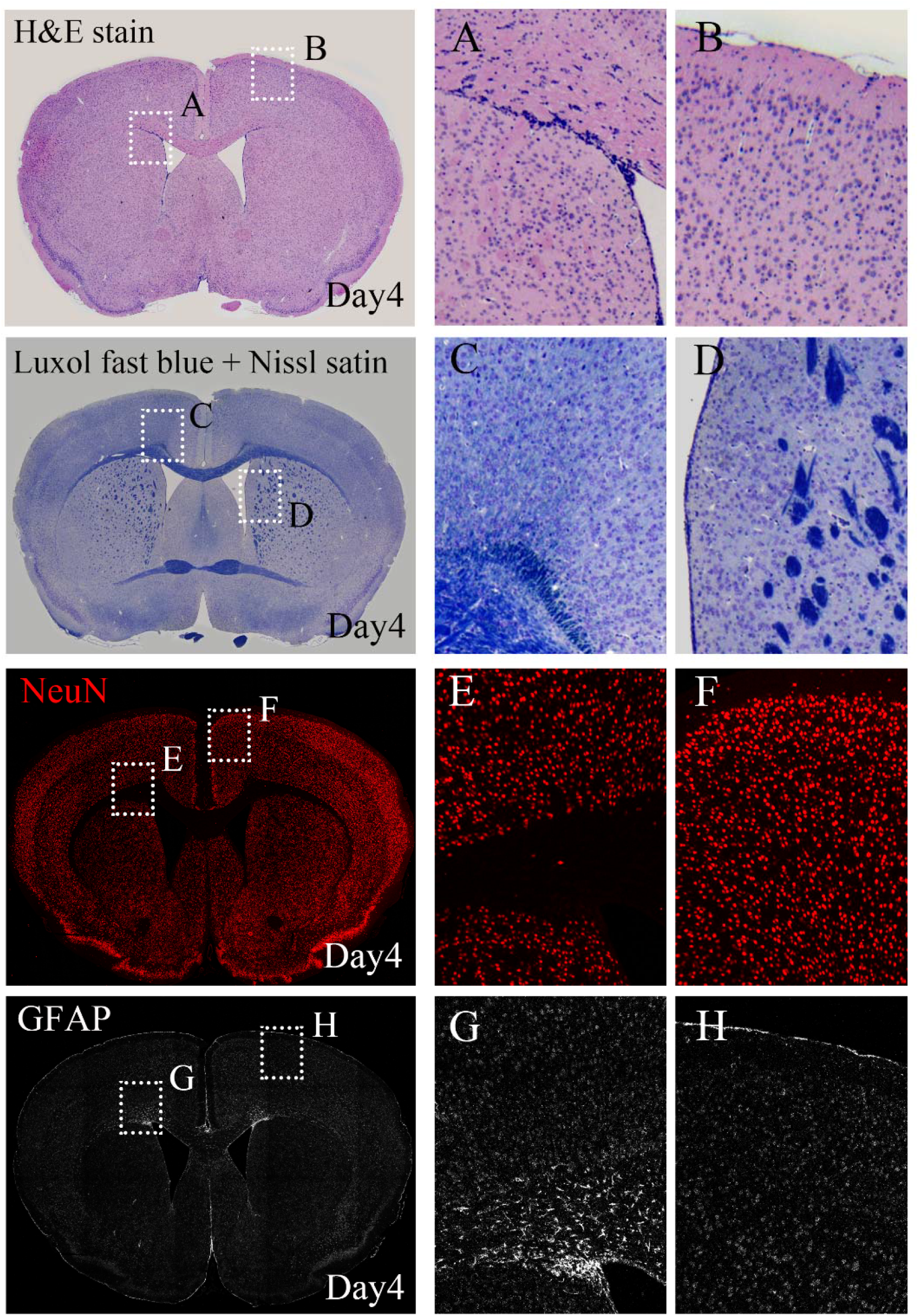
Safety evaluation of VLIUS stimulation at day 4. (A, B) Hematoxylin and Eosin stain: no significant tissue changes. (C, D) Luxol fast blue and Nissl stain: no significant myelin sheath changes or decrease in neuronal cells. (E, F) Neuronal nuclei staining: no decrease in mature neurons. (G, H) GFAP staining: no increase of astrocytes, indicating no neuroinflammation.

## References and Notes

1. J. J. Iliff, M. Wang, Y. Liao, B. A. Plogg, W. Peng, G. A. Gundersen, H. Benveniste, G. E. Vates, R. Deane, S. A. Goldman, E. A. Nagelhus, M. Nedergaard, A paravascular pathway facilitates CSF flow through the brain parenchyma and the clearance of interstitial solutes, including amyloid beta. Sci Transl Med 4, 147ra111 (2012).

2. J. J. Iliff, H. Lee, M. Yu, T. Feng, J. Logan, M. Nedergaard, H. Benveniste, Brain-wide pathway for waste clearance captured by contrast-enhanced MRI. J Clin Invest 123, 1299–1309 (2013).

3. M. Nedergaard, Neuroscience. Garbage truck of the brain. Science 340, 1529–1530 (2013).

4. L. Xie, H. Kang, Q. Xu, M. J. Chen, Y. Liao, M. Thiyagarajan, J. O’Donnell, D. J. Christensen, C. Nicholson, J. J. Iliff, T. Takano, R. Deane, M. Nedergaard, Sleep drives metabolite clearance from the adult brain. Science 342, 373–377 (2013).

5. B. A. Plog, M. L. Dashnaw, E. Hitomi, W. Peng, Y. Liao, N. Lou, R. Deane, M. Nedergaard, Biomarkers of traumatic injury are transported from brain to blood via the glymphatic system. J Neurosci 35, 518–526 (2015).

6. W. Peng, T. M. Achariyar, B. Li, Y. Liao, H. Mestre, E. Hitomi, S. Regan, T. Kasper, S. Peng, F. Ding, H. Benveniste, M. Nedergaard, R. Deane, Suppression of glymphatic fluid transport in a mouse model of Alzheimer’s disease. Neurobiol Dis 93, 215–225 (2016).

7. B. C. Reeves, J. K. Karimy, A. J. Kundishora, H. Mestre, H. M. Cerci, C. Matouk, S. L. Alper, I. Lundgaard, M. Nedergaard, K. T. Kahle, Glymphatic System Impairment in Alzheimer’s Disease and Idiopathic Normal Pressure Hydrocephalus. Trends Mol Med 26, 285–295 (2020).

8. J. J. Iliff, M. J. Chen, B. A. Plog, D. M. Zeppenfeld, M. Soltero, L. Yang, I. Singh, R. Deane, M. Nedergaard, Impairment of glymphatic pathway function promotes tau pathology after traumatic brain injury. J Neurosci 34, 16180–16193 (2014).

9. M. J. Sullan, B. M. Asken, M. S. Jaffee, S. T. DeKosky, R. M. Bauer, Glymphatic system disruption as a mediator of brain trauma and chronic traumatic encephalopathy. Neurosci Biobehav Rev 84, 316–324 (2018).

10. H. Mestre, T. Du, A. M. Sweeney, G. Liu, A. J. Samson, W. Peng, K. N. Mortensen, F. F. Staeger, P. A. R. Bork, L. Bashford, E. R. Toro, J. Tithof, D. H. Kelley, J. H. Thomas, P. G. Hjorth, E. A. Martens, R. I. Mehta, O. Solis, P. Blinder, D. Kleinfeld, H. Hirase, Y. Mori, M. Nedergaard, Cerebrospinal fluid influx drives acute ischemic tissue swelling. Science 367, (2020).

11. R. Goulay, J. Flament, M. Gauberti, M. Naveau, N. Pasquet, C. Gakuba, E. Emery, P. Hantraye, D. Vivien, R. Aron-Badin, T. Gaberel, Subarachnoid Hemorrhage Severely Impairs Brain Parenchymal Cerebrospinal Fluid Circulation in Nonhuman Primate. Stroke 48, 2301–2305 (2017).

12. A. J. Schain, A. Melo-Carrillo, A. M. Strassman, R. Burstein, Cortical Spreading Depression Closes Paravascular Space and Impairs Glymphatic Flow: Implications for Migraine Headache. J Neurosci 37, 2904–2915 (2017).

13. A. Carotenuto, L. Cacciaguerra, E. Pagani, P. Preziosa, M. Filippi, M. A. Rocca, Glymphatic system impairment in multiple sclerosis: relation with brain damage and disability. Brain, (2021).

14. A. Zamani, A. K. Walker, B. Rollo, K. L. Ayers, R. Farah, T. J. O’Brien, D. K. Wright, Impaired glymphatic function in the early stages of disease in a TDP-43 mouse model of amyotrophic lateral sclerosis. Transl Neurodegener 11, 17 (2022).

15. T. M. Achariyar, B. Li, W. Peng, P. B. Verghese, Y. Shi, E. McConnell, A. Benraiss, T. Kasper, W. Song, T. Takano, D. M. Holtzman, M. Nedergaard, R. Deane, Glymphatic distribution of CSF-derived apoE into brain is isoform specific and suppressed during sleep deprivation. Mol Neurodegener 11, 74 (2016).

16. H. Lee, L. Xie, M. Yu, H. Kang, T. Feng, R. Deane, J. Logan, M. Nedergaard, H. Benveniste, The Effect of Body Posture on Brain Glymphatic Transport. J Neurosci 35, 11034–11044 (2015).

17. H. Mestre, J. Tithof, T. Du, W. Song, W. Peng, A. M. Sweeney, G. Olveda, J. H. Thomas, M. Nedergaard, D. H. Kelley, Flow of cerebrospinal fluid is driven by arterial pulsations and is reduced in hypertension. Nat Commun 9, 4878 (2018).

18. B. T. Kress, J. J. Iliff, M. Xia, M. Wang, H. S. Wei, D. Zeppenfeld, L. Xie, H. Kang, Q. Xu, J. A. Liew, B. A. Plog, F. Ding, R. Deane, M. Nedergaard, Impairment of paravascular clearance pathways in the aging brain. Ann Neurol 76, 845–861 (2014).

19. C. Gakuba, T. Gaberel, S. Goursaud, J. Bourges, C. Di Palma, A. Quenault, S. Martinez de Lizarrondo, D. Vivien, M. Gauberti, General Anesthesia Inhibits the Activity of the “Glymphatic System”. Theranostics 8, 710–722 (2018).

20. L. M. Hablitz, H. S. Vinitsky, Q. Sun, F. F. Staeger, B. Sigurdsson, K. N. Mortensen, T.. Lilius, M. Nedergaard, Increased glymphatic influx is correlated with high EEG delta power and low heart rate in mice under anesthesia. Sci Adv 5, eaav5447 (2019).

21. H. Mestre, L. M. Hablitz, A. L. Xavier, W. Feng, W. Zou, T. Pu, H. Monai, G. Murlidharan, R. M. Castellanos Rivera, M. J. Simon, M. M. Pike, V. Pla, T. Du, B. T. Kress, X. Wang, B. A. Plog, A. S. Thrane, I. Lundgaard, Y. Abe, M. Yasui, J. H. Thomas, M. Xiao, H. Hirase, A. Asokan, J. J. Iliff, M. Nedergaard, Aquaporin-4-dependent glymphatic solute transport in the rodent brain. Elife 7, (2018).

22. P. Kitchen, M. M. Salman, A. M. Halsey, C. Clarke-Bland, J. A. MacDonald, H. Ishida, H. J. Vogel, S. Almutiri, A. Logan, S. Kreida, T. Al-Jubair, J. Winkel Missel, P. Gourdon, S. Tornroth-Horsefield, M. T. Conner, Z. Ahmed, A. C. Conner, R. M. Bill, Targeting Aquaporin-4 Subcellular Localization to Treat Central Nervous System Edema. Cell 181, 784–799 e719 (2020).

23. V. Benfenati, M. Caprini, M. Dovizio, M. N. Mylonakou, S. Ferroni, O. P. Ottersen, M. Amiry-Moghaddam, An aquaporin-4/transient receptor potential vanilloid 4 (AQP4/TRPV4) complex is essential for cell-volume control in astrocytes. Proc Natl Acad Sci U S A 108, 2563–2568 (2011).

24. M. M. Maneshi, B. Maki, R. Gnanasambandam, S. Belin, G. K. Popescu, F. Sachs, S. Z. Hua, Mechanical stress activates NMDA receptors in the absence of agonists. Sci Rep 7, 39610 (2017).

25. W.-H. Liao, M.-Y. Hsiao, Y. Kung, H.-L. Liu, J.-C. Béra, C. Inserra, W.-S. Chen, TRPV4 promotes acoustic wave-mediated BBB opening via Ca2+/PKC-δ pathway. Journal of Advanced Research 26, 15–28 (2020).

26. M. Aryal, M. M. Azadian, A. R. Hart, N. Macedo, Q. Zhou, E. L. Rosenthal, R. D. Airan, Noninvasive ultrasonic induction of cerebrospinal fluid flow enhances intrathecal drug delivery. J Control Release 349, 434–442 (2022).

27. Y. Lee, Y. Choi, E. J. Park, S. Kwon, H. Kim, J. Y. Lee, D. S. Lee, Improvement of glymphatic-lymphatic drainage of beta-amyloid by focused ultrasound in Alzheimer’s disease model. Sci Rep 10, 16144 (2020).

28. J. Lim, H. H. Tai, W. H. Liao, Y. C. Chu, C. M. Hao, Y. C. Huang, C. H. Lee, S. S. Lin, S. Hsu, Y. C. Chien, D. M. Lai, W. S. Chen, C. C. Chen, J. L. Wang, ASIC1a is required for neuronal activation via low-intensity ultrasound stimulation in mouse brain. Elife 10, (2021).

29. Y. C. Chu, J. Lim, C. H. Lai, M. C. Tseng, Y. S. Chu, J. L. Wang, Elevation of Intra-Cellular Calcium in Nucleus Pulposus Cells with Micro-Pipette-Guided Ultrasound. Ultrasound Med Biol 47, 1775–1784 (2021).

30. O. Butenko, D. Dzamba, J. Benesova, P. Honsa, V. Benfenati, V. Rusnakova, S. Ferroni, M. Anderova, The increased activity of TRPV4 channel in the astrocytes of the adult rat hippocampus after cerebral hypoxia/ischemia. PLoS One 7, e39959 (2012).

31. C. Rakers, M. Schmid, G. C. Petzold, TRPV4 channels contribute to calcium transients in astrocytes and neurons during peri-infarct depolarizations in a stroke model. Glia 65, 1550–1561 (2017).

32. K. M. Dunn, D. C. Hill-Eubanks, W. B. Liedtke, M. T. Nelson, TRPV4 channels stimulate Ca2+-induced Ca2+ release in astrocytic endfeet and amplify neurovascular coupling responses. Proc Natl Acad Sci U S A 110, 6157–6162 (2013).

33. M. Saifeddine, M. El-Daly, K. Mihara, N. W. Bunnett, P. McIntyre, C. Altier, M. D. Hollenberg, R. Ramachandran, GPCR-mediated EGF receptor transactivation regulates TRPV4 action in the vasculature. Br J Pharmacol 172, 2493–2506 (2015).

34. W. G. Darby, S. Potocnik, R. Ramachandran, M. D. Hollenberg, O. L. Woodman, P. McIntyre, Shear stress sensitizes TRPV4 in endothelium-dependent vasodilatation. Pharmacol Res 133, 152–159 (2018).

35. J. J. Iliff, M. Wang, D. M. Zeppenfeld, A. Venkataraman, B. A. Plog, Y. Liao, R. Deane, M. Nedergaard, Cerebral arterial pulsation drives paravascular CSF-interstitial fluid exchange in the murine brain. J Neurosci 33, 18190–18199 (2013).

36. P. Rajasekhar, D. P. Poole, N. A. Veldhuis, Role of Nonneuronal TRPV4 Signaling in Inflammatory Processes. Adv Pharmacol 79, 117–139 (2017).

37. H. Mestre, Y. Mori, M. Nedergaard, The Brain’s Glymphatic System: Current Controversies. Trends Neurosci, (2020).

38. A. O. Jo, D. A. Ryskamp, T. T. Phuong, A. S. Verkman, O. Yarishkin, N. MacAulay, D. Krizaj, TRPV4 and AQP4 Channels Synergistically Regulate Cell Volume and Calcium Homeostasis in Retinal Muller Glia. J Neurosci 35, 13525–13537 (2015).

39. Y. Takayama, K. Shibasaki, Y. Suzuki, A. Yamanaka, M. Tominaga, Modulation of water efflux through functional interaction between TRPV4 and TMEM16A/anoctamin 1. FASEB J 28, 2238–2248 (2014).

40. K. Shibasaki, TRPV4 activation by thermal and mechanical stimuli in disease progression. Lab Invest 100, 218–223 (2020).

41. A. Iuso, D. Krizaj, TRPV4-AQP4 interactions ‘turbocharge’ astroglial sensitivity to small osmotic gradients. Channels (Austin) 10, 172–174 (2016).

42. M. G. Mola, E. Saracino, F. Formaggio, A. G. Amerotti, B. Barile, T. Posati, A. Cibelli, A. Frigeri, C. Palazzo, R. Zamboni, M. Caprini, G. P. Nicchia, V. Benfenati, Cell Volume Regulation Mechanisms in Differentiated Astrocytes. Cell Physiol Biochem 55, 196–212 (2021).

43. M. M. Salman, P. Kitchen, M. N. Woodroofe, J. E. Brown, R. M. Bill, A. C. Conner, M. T. Conner, Hypothermia increases aquaporin 4 (AQP4) plasma membrane abundance in human primary cortical astrocytes via a calcium/transient receptor potential vanilloid 4 (TRPV4)- and calmodulin-mediated mechanism. Eur J Neurosci 46, 2542–2547 (2017).

44. J. Jorgacevski, R. Zorec, M. Potokar, Insights into Cell Surface Expression, Supramolecular Organization, and Functions of Aquaporin 4 Isoforms in Astrocytes. Cells 9, (2020).

45. M. G. Mola, A. Sparaneo, C. D. Gargano, D. C. Spray, M. Svelto, A. Frigeri, E. Scemes, G. P. Nicchia, The speed of swelling kinetics modulates cell volume regulation and calcium signaling in astrocytes: A different point of view on the role of aquaporins. Glia 64, 139–154 (2016).

46. J. Liu, Y. Yang, X. Li, P. Zhang, Y. Qi, H. Hu, in Methods in Enzymology, M. Fukuda, Ed. (Academic Press, 2010), vol. 479, pp. 353–366.

47. D. Becker, C. Blase, J. Bereiter-Hahn, M. Jendrach, TRPV4 exhibits a functional role in cell-volume regulation. J Cell Sci 118, 2435–2440 (2005).

48. V. Benfenati, M. Amiry-Moghaddam, M. Caprini, M. N. Mylonakou, C. Rapisarda, O. P. Ottersen, S. Ferroni, Expression and functional characterization of transient receptor potential vanilloid-related channel 4 (TRPV4) in rat cortical astrocytes. Neuroscience 148, 876–892 (2007).

49. T. J. Lohela, T. O. Lilius, M. Nedergaard, The glymphatic system: implications for drugs for central nervous system diseases. Nat Rev Drug Discov 21, 763–779 (2022).

50. S. Xu, D. Ye, L. Wan, Y. Shentu, Y. Yue, M. Wan, H. Chen, Correlation Between Brain Tissue Damage and Inertial Cavitation Dose Quantified Using Passive Cavitation Imaging. Ultrasound Med Biol 45, 2758–2766 (2019).

51. D. G. Blackmore, F. Turpin, T. Palliyaguru, H. T. Evans, A. Chicoteau, W. Lee, M. Pelekanos, N. Nguyen, J. Song, R. K. P. Sullivan, P. Sah, P. F. Bartlett, J. Gotz, Low-intensity ultrasound restores long-term potentiation and memory in senescent mice through pleiotropic mechanisms including NMDAR signaling. Mol Psychiatry, (2021).

52. J. A. Stokum, M. S. Kwon, S. K. Woo, O. Tsymbalyuk, R. Vennekens, V. Gerzanich, J. M. Simard, SUR1-TRPM4 and AQP4 form a heteromultimeric complex that amplifies ion/water osmotic coupling and drives astrocyte swelling. Glia 66, 108–125 (2018).

53. B. Reiter, R. Kraft, D. Gunzel, S. Zeissig, J. D. Schulzke, M. Fromm, C. Harteneck, TRPV4-mediated regulation of epithelial permeability. FASEB J 20, 1802–1812 (2006).

54. H. Zhao, K. Zhang, R. Tang, H. Meng, Y. Zou, P. Wu, R. Hu, X. Liu, H. Feng, Y. Chen, TRPV4 Blockade Preserves the Blood-Brain Barrier by Inhibiting Stress Fiber Formation in a Rat Model of Intracerebral Hemorrhage. Front Mol Neurosci 11, 97 (2018).

55. P. Jie, Z. Lu, Z. Hong, L. Li, L. Zhou, Y. Li, R. Zhou, Y. Zhou, Y. Du, L. Chen, L. Chen, Activation of Transient Receptor Potential Vanilloid 4 is Involved in Neuronal Injury in Middle Cerebral Artery Occlusion in Mice. Mol Neurobiol 53, 8–17 (2016).

56. P. Jie, Z. Hong, Y. Tian, Y. Li, L. Lin, L. Zhou, Y. Du, L. Chen, L. Chen, Activation of transient receptor potential vanilloid 4 induces apoptosis in hippocampus through downregulating PI3K/Akt and upregulating p38 MAPK signaling pathways. Cell Death Dis 6, e1775 (2015).

57. D. Guo, J. Zou, N. Rensing, M. Wong, In Vivo Two-Photon Imaging of Astrocytes in GFAP-GFP Transgenic Mice. PLoS One 12, e0170005 (2017).

58. V. Benfenati, G. P. Nicchia, M. Svelto, C. Rapisarda, A. Frigeri, S. Ferroni, Functional down-regulation of volume-regulated anion channels in AQP4 knockdown cultured rat cortical astrocytes. J Neurochem 100, 87–104 (2007).

59. S. Baratchi, P. Keov, W. G. Darby, A. Lai, K. Khoshmanesh, P. Thurgood, P. Vahidi, K. Ejendal, P. McIntyre, The TRPV4 Agonist GSK1016790A Regulates the Membrane Expression of TRPV4 Channels. Front Pharmacol 10, 6 (2019).

60. A. Imura, Y. Tsuji, M. Murata, R. Maeda, K. Kubota, A. Iwano, C. Obuse, K. Togashi, M. Tominaga, N. Kita, K. Tomiyama, J. Iijima, Y. Nabeshima, M. Fujioka, R. Asato, S. Tanaka, K. Kojima, J. Ito, K. Nozaki, N. Hashimoto, T. Ito, T. Nishio, T. Uchiyama, T. Fujimori, Y. Nabeshima, alpha-Klotho as a regulator of calcium homeostasis. Science 316, 1615–1618 (2007).

61. J. Lim, Y.-C. Chu, C.-M. Hao, W.-H. Liao, S.-S. Lin, S. Hsu, H.-H. Tai, Y.-C. Chien, D.-M. Lai, W.-S. Chen, C.-C. Chen, J.-L. Wang, ASIC1a is required for neuronal activation via low-intensity ultrasound stimulation in mouse brain. bioRxiv, 2020.2007.2010.196634 (2020).

62. A. L. R. Xavier, N. L. Hauglund, S. von Holstein-Rathlou, Q. Li, S. Sanggaard, N. Lou, I. Lundgaard, M. Nedergaard, Cannula Implantation into the Cisterna Magna of Rodents. J Vis Exp, (2018).

63. F. Galland, M. Seady, J. Taday, S. S. Smaili, C. A. Goncalves, M. C. Leite, Astrocyte culture models: Molecular and function characterization of primary culture, immortalized astrocytes and C6 glioma cells. Neurochem Int 131, 104538 (2019).

