## Supplementary figures and images for "Enhancing glymphatic function with very low-intensity ultrasound via the transient receptor potential vanilloid-4-aquaporin-4 pathway"

### Supplementary Figure 1

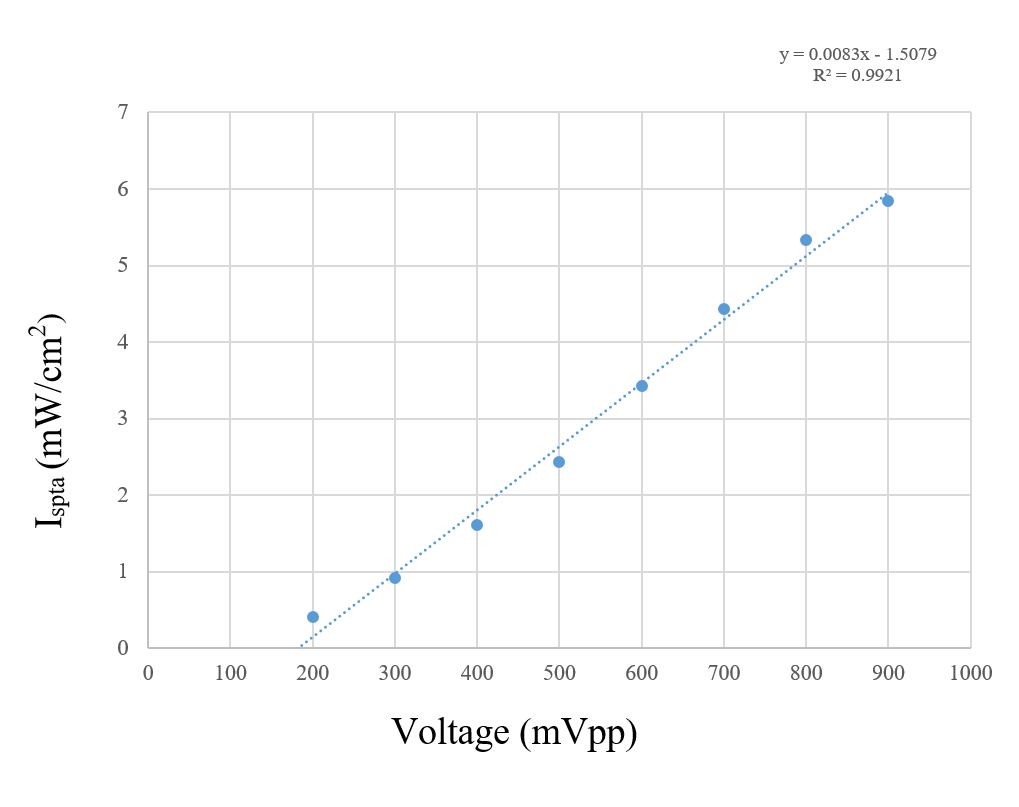

### Supplementary Figure 2

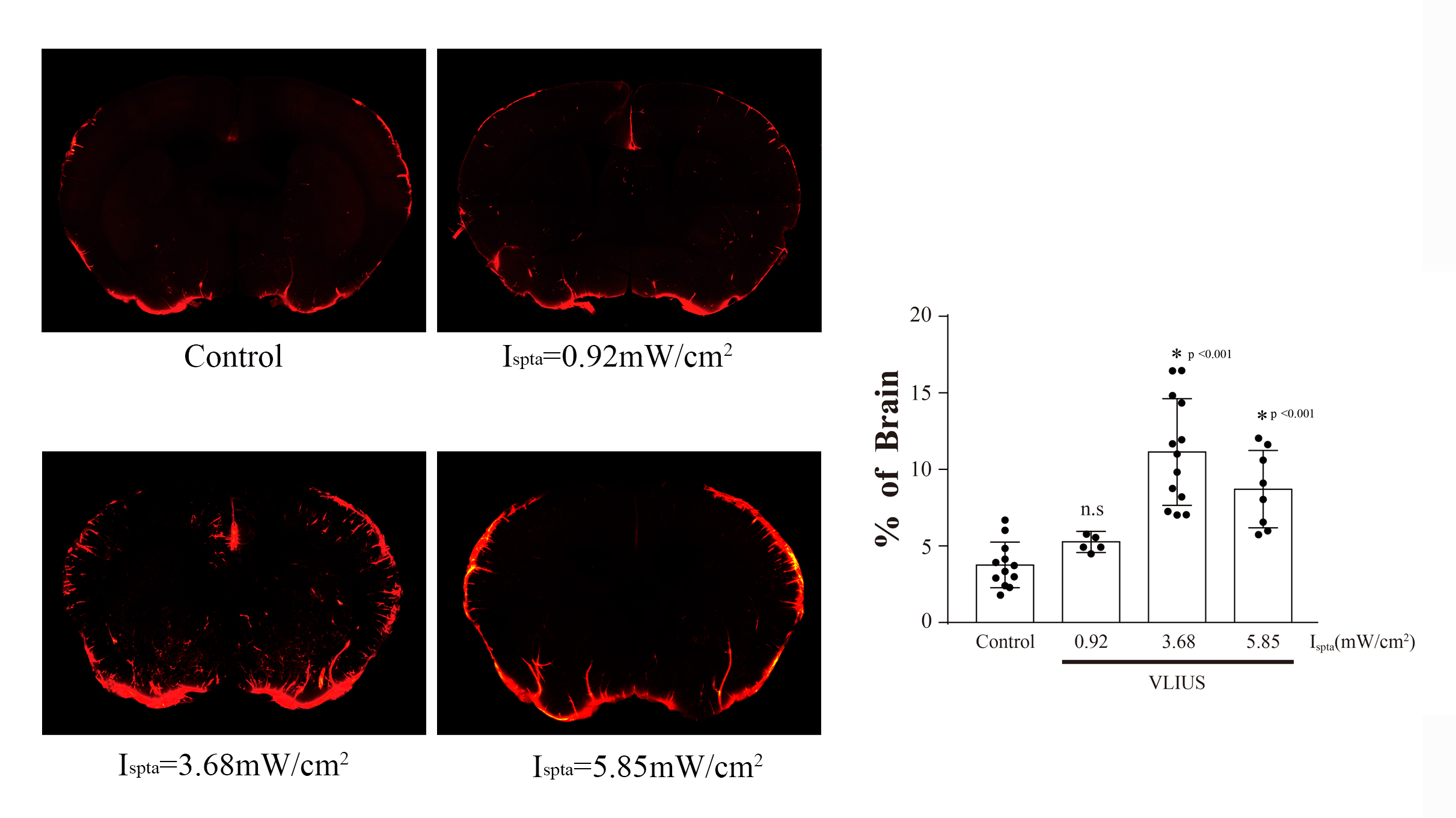

### Supplementary Figure 3

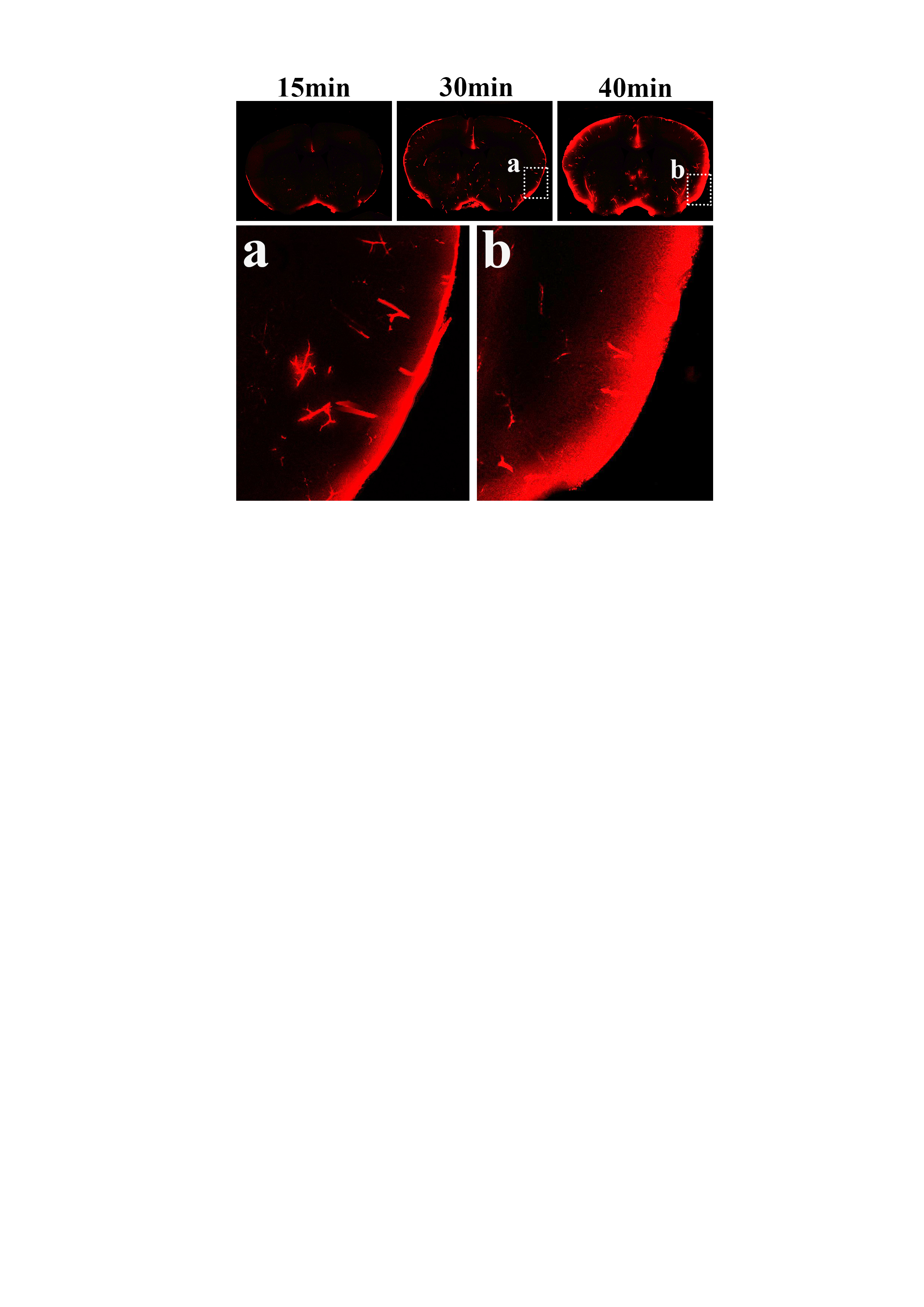

### Supplementary Figure 4

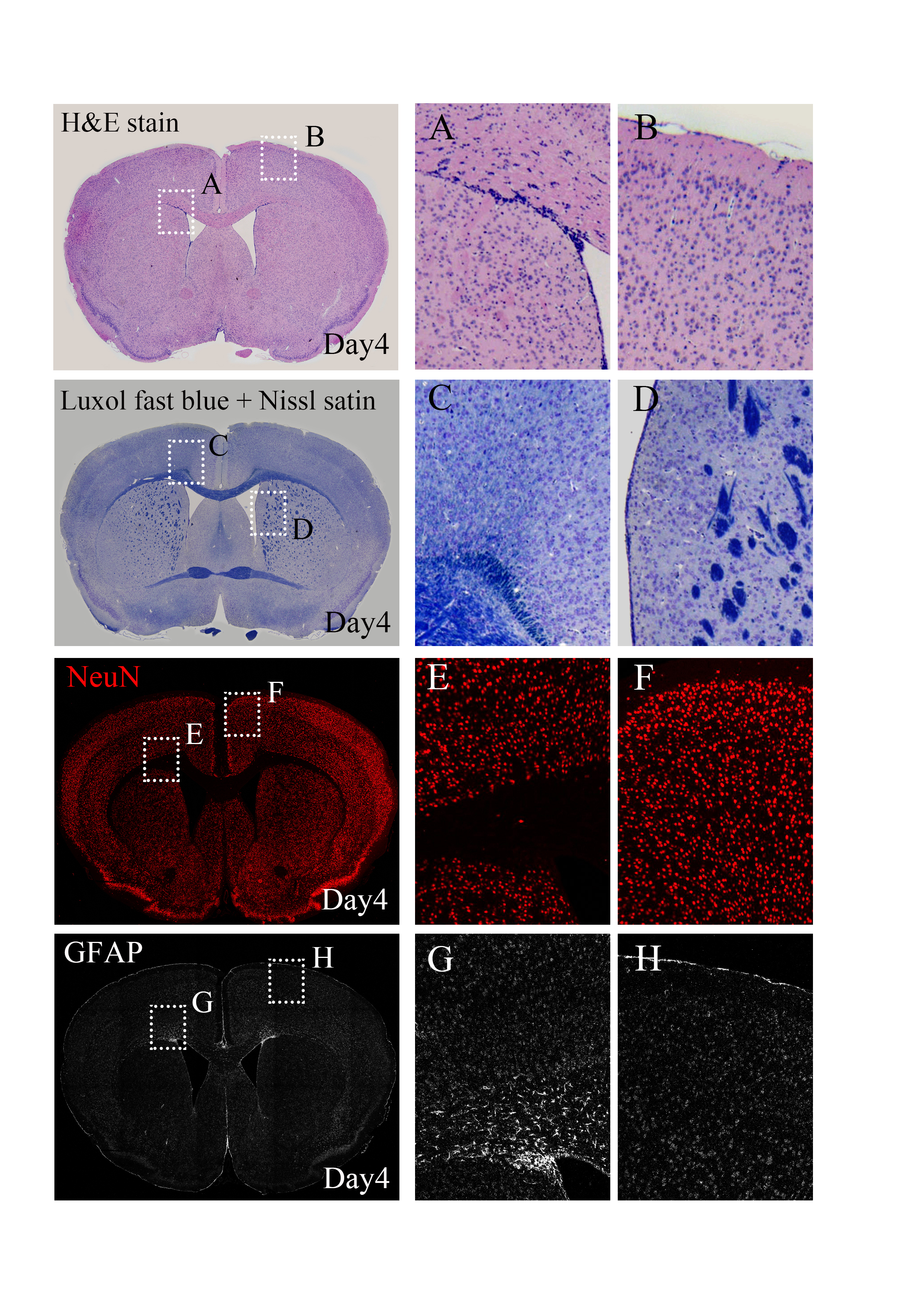
